# Aiki-XP: leakage-controlled multimodal prediction of within-species relative protein expression at pan-bacterial scale

**DOI:** 10.64898/2026.04.21.719525

**Authors:** Hudson Tien, Radheesh Sharma Meda, Shankar Shastry, Venkatesh Mysore

## Abstract

Generalizable protein-expression prediction can accelerate protein engineering, inform disease mechanisms, and help optimize heterologous recombinant protein production. Protein expression is governed by many interacting parameters that no single omics view captures. We develop Aiki-XP, a multimodal platform integrating four biological scales (genome, operon, coding sequence, protein) plus biophysical features across 492,026 genes from 385 bacterial species. Aiki-XP predicts within-species relative abundance (per-species z-score rank), not absolute copies per cell. Under a leakage-controlled gene-operon split Aiki-XP reaches Spearman ***ρ*_nc_ = 0.592** versus **0.509** for ESM-C 600M alone, and each tier of a monotone protein**→**operon**→**genome deployment ladder yields a statistically significant gain; a five-recipe rank-average ensemble adds a further **+0.016**. All recipes were locked before external evaluation; transfer to heterologous, cross-species, and novel-phylum benchmarks demonstrates utility and limits. Ablations and scaling experiments identify operon-scale genomic context, not protein-language-model capacity, as the rate-limiting input at this scale; one foundation model per biological scale suffices, with same-scale stacking adding little.

## Introduction

Protein expression is an essential feature of life, required for growth, maintenance, and repair [42]. Understanding how genetic information is converted into protein molecules is critical for understanding the mechanisms underpinning biological processes and consequently their role in disease biology. Further, engineered proteins produced in bacterial hosts have become a mainstay of therapeutics, have manifold industrial uses, and function as biomaterials in many real-world applications [43]. In this context, models that accurately predict protein expression enable *in silico* screening of protein variants before experimental testing, saving substantial time and resources [33, 41].

Protein expression in prokaryotes was first described by the operon model [44] and is governed by multiple parameters including genetic information, operon architecture, transcription, RNA stability, translation, and growth-law constraints [28]. Despite growing interest in computational expression prediction, no existing approach integrates all of these regulatory layers at scale and evaluates cross-species generalization under stringent homology control.

The building blocks for such integration now exist: foundation models provide rich learned representations for proteins[7, 35], coding and operon-level DNA[8, 12], and a genome-contextualised view of each protein in its native chromosomal neighborhood[13]; recent work has begun fusing two or three of these for expression-related tasks[29, 32, 36, 39]. Two fundamental challenges, however, have kept this potential from being realized at pan-bacterial scale. Curating expression data across hundreds of species requires linking expression measurements to their genomic and regulatory context across multiple databases, a pipeline complexity that has prevented any large-scale benchmark from being assembled. Even where benchmarks exist, conserved gene families such as ribosomal proteins and elongation factors span nearly all bacteria and dominate standard train/test splits, so that models are rewarded for recognizing families rather than learning regulatory signals[1, 2]. As a result, fusion efforts to date have been limited to ≤7K sequences from single species[38], leaving open whether multi-scale integration genuinely improves prediction or relies on recognizing conserved families. Resolving this question requires both a dataset large enough to absorb the statistical cost of strict family-level holdout and a modeling framework that tests each biological scale independently.

Here we present Aiki-XP, a multimodal fusion platform addressing both gaps across 492,026 genes from 385 bacterial species. Aiki-XP (the platform) instantiates a monotone deployment ladder of five input tiers; XP5 denotes the 5-modality top tier used for the in-distribution benchmark throughout this paper. An operon in this paper refers to a co-transcribed cluster of functionally related bacterial genes sharing a single mRNA; the conserved “ribosomal component” is a single connected sequence-based gene cluster covering ribosomal proteins and tightly linked partners that reaches every species in our dataset. Aiki-XP predicts within-species relative expression on a per-species z-score scale derived from the multi-study integrated abundances in PaXDb[4] and the intensity-based absolute-quantification (iBAQ) atlas of Abele 2025[5]: each prediction is a within-organism rank. This rank-based target is invariant to per-species linear rescaling across quantification platforms, and aligns learning with the regulatory features (codon adaptation, RNA folding, operon architecture) that bacterial expression actually depends on. We use this normalized endpoint to separate two non-equivalent generalization tasks that standard benchmarks conflate: prediction for *novel gene families* within known species, and transfer to *novel species* where many known families remain available. A model can appear strong under one regime while remaining weak under the other; the pervasive random split characterizes neither real-world scenario.

## Results

### A 492K-gene, 385-species benchmark with homology-controlled splits

We collated expression measurements for 492,026 genes across 385 bacterial species spanning five phyla: Pseudomonadota (162 species), Bacillota (162), Actinomycetota (27), Bacteroidota (9), and 25 species from minor phyla including Cyanobacteriota, Deinococcota, Campylobacterota, and Spirochaetota (Fig. 1a). Data were integrated from multi-study proteomics via PaXDb v6.0[4] (∼235K genes) and single-study iBAQ measurements from the bacterial proteome atlas of Abele et al. 2025[5] (256K genes, hereafter “Abele”). Assembling this benchmark required resolving gene↔protein↔genome mappings, operon annotations, and coding-sequence (CDS) boundaries across all 385 genomes, then extracting embeddings from 12 foundation models, with the 492K set surviving the pipeline that had to discard ∼40% of candidate genes due to incomplete annotations or unresolvable cross-database identifiers (see Online Methods).

**Fig. 1.**
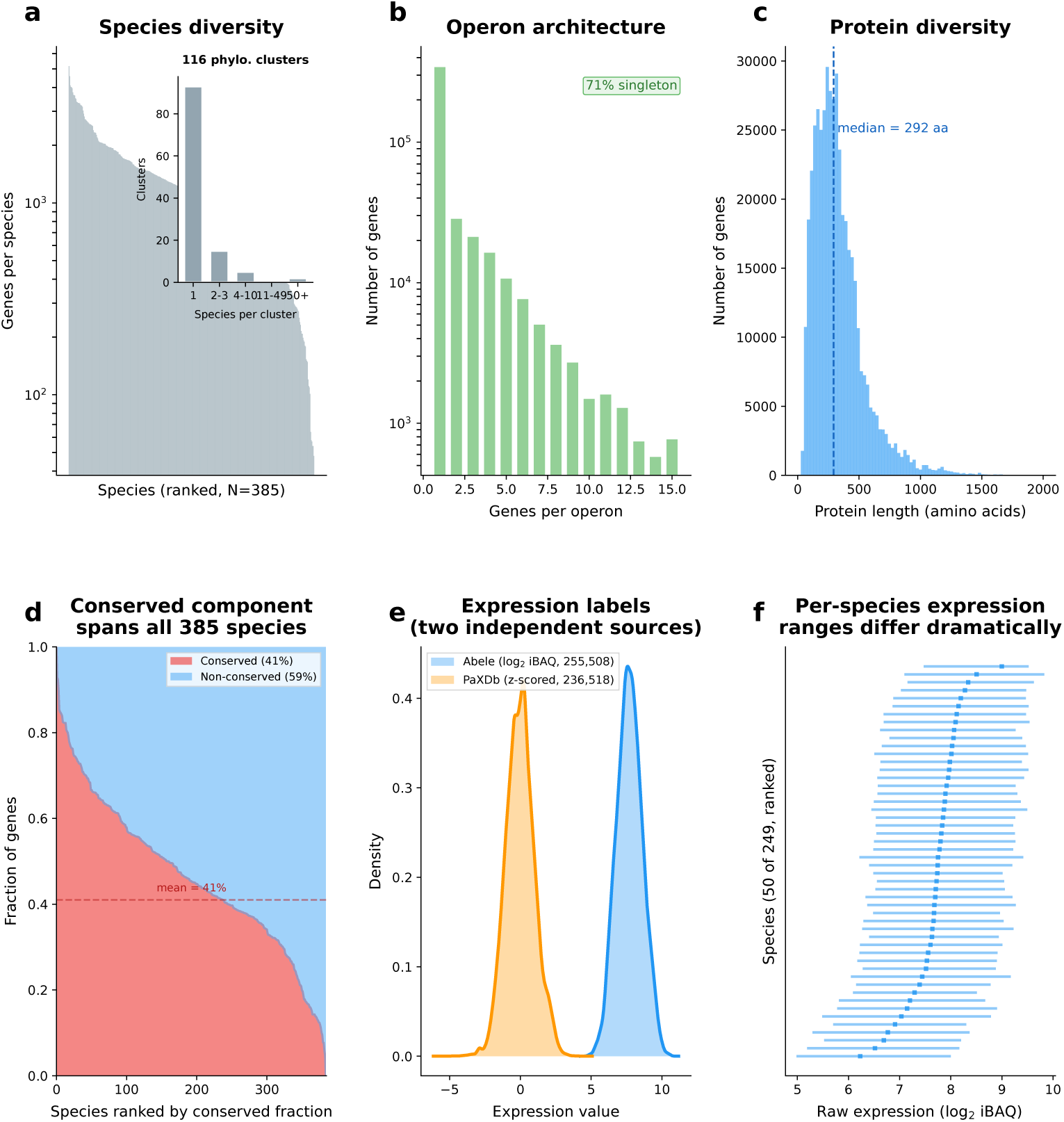
A pan-bacterial expression benchmark: 492,026 genes across 385 species. **a,** Species diversity: genes per species on log scale, with a long tail from model organisms to environmental bacteria. Inset: species group into 116 phylogenetic clusters at the Mash threshold used for species-cluster evaluation (*t* = 0.20). **b,** Operon architecture: 71% of genes are singletons; multi-gene operons reach 15+ genes. Multi-gene operons in this dataset are assigned a single per-operon expression value reflecting the proteomics measurement aggregation (both PaXDb and Abele sources integrate at the operon/protein-group level); the leak-free test of multimodal fusion is therefore on singletons, where every gene carries a unique label. **c,** Protein length distribution (median 292 amino acids). **d,** The conserved ribosomal component: 202,945 genes (41%) form a single connected component spanning all 385 species. We report the Spearman rank correlation coefficient *ρ*nc computed over the non-conserved genes (59% of the dataset) as the primary metric throughout. **e,** Raw expression labels from two independent proteomics sources: PaXDb (multi-study integrated, ∼235K genes, reported in ppm and as z-scored per species) and the Abele atlas (single-study iBAQ, 256K genes, log_2_ iBAQ). The model predicts per-species z-scored expression, not raw values (Supplementary Fig. S2). **f,** Per-species raw expression ranges for Abele species (50 of 249, log_2_ iBAQ): means span ∼5 orders of magnitude and within-species ranges vary 3–6 log_2_ units, motivating per-species z-score normalization. PaXDb values are delivered pre-normalized; see Supplementary Fig. S2 for the z-scoring justification. The phylogenetic relationships across the 385 species _1_a_4_re shown in Supplementary Fig. S7.

Homology leakage is a central challenge for expression benchmarks[1–3]: conserved gene families that occur in both training and test sets inflate apparent performance. Our gene-operon split ensures there is no cross-over within the 129,078 MMseqs2[18] protein clusters at 50% identity. A bipartite graph analysis revealed that 202,945 genes (41%) that were predominantly ribosomal proteins and associated operon partners[6] form a single conserved component spanning all 385 species (Fig. 1d). This operon cluster alone was proportionally distributed across the splits to enable monitoring the effect of data leakage without affecting the non-conserved genes (Extended Data Fig. ED2). Since these conserved genes are substantially easier to predict than the remaining 59%, we report performance on non-conserved genes (*ρ*_nc_) as the primary metric throughout, alongside overall *ρ* for comparability with prior work.

### The XP5 architecture: four biological scales and a biophysical block

The multimodal model (XP5) integrates four biological scales, each captured by a dedicated foundation model (Fig. 2a; Extended Data Table ED3):

- **Genome context** (Bacformer-large, 960d): a recently-introduced transformer[13] that ingests an ESM-2 protein embedding *together with* its chromosomal neighbors, producing a per-gene representation conditioned on its position in the host genome; the embedding therefore carries information about operon order, chromosomal regulation, and gene-neighborhood composition that protein-sequence models alone cannot access.
- **Operon architecture** (Evo-2 7B, 4,096d): full polycistronic transcription unit including intergenic regions.
- **Coding-sequence composition** (HyenaDNA, 256d): codon usage, GC structure, and CDS-level regulatory signals.
- **Protein identity** (ESM-C, 1,152d): amino acid sequence and evolutionary conservation.

**Fig. 2.**
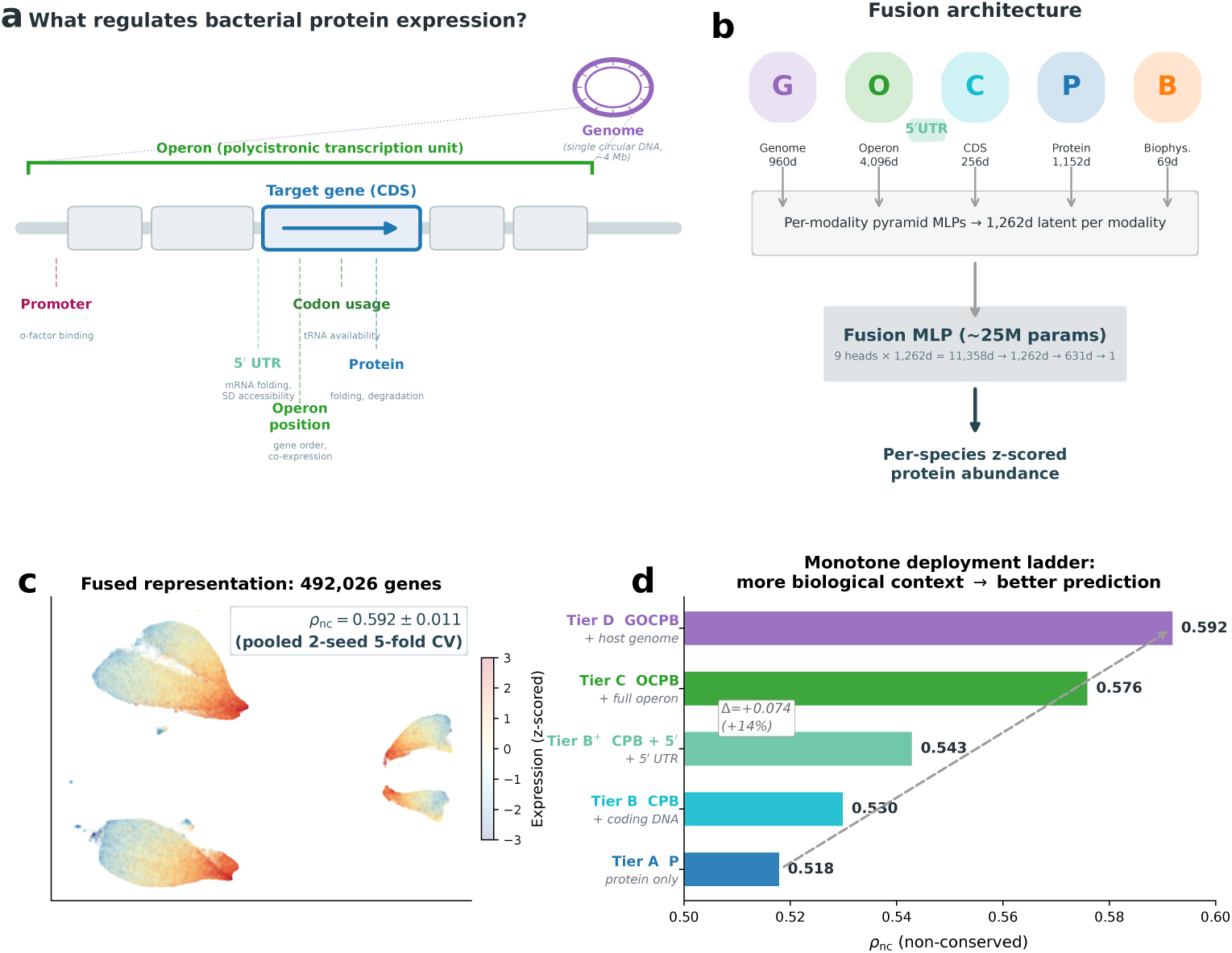
The Aiki-XP platform: four biological scales with one foundation model each after selection. **a,** Four nested biological scales govern expression: genome context, operon architecture, coding-sequence composition, and protein identity, each captured by a dedicated foundation model in the selected XP5 recipe. Per-gene biophysical features (69 dimensions) bridge all scales with mechanistic signals that operon-level mean-pooling discards: translation-initiation thermodynamics, codon adaptation, operon position, protein physicochemistry, and intrinsic disorder. Twelve foundation models were evaluated; the four shown here plus the biophysical block were selected by greedy forward stepwise on *ρ*nc. ProtT5-XL (3B; 1,024-dimensional embeddings)[^35^] is used as a second protein-level representation in protein-only Tier A (see text). **b,** Per-modality pyramid adapters project each embedding onto a common 1,262-dimensional latent space (five modalities: genome 960d, operon 4,096d, CDS 256d, protein 1,152d, and a biophysical-features block 69d; 6,533 input dimensions in all). The adapter outputs are concatenated and fed to a fusion multi-layer perceptron (1,262d → 631d → 1 scalar; ∼25M parameters total). **c,** UMAP of the 1,262d penultimate fusion representation (492,026 genes, colored by z-scored expression). The multi-scale fusion produces a smooth expression gradient that single-scale inputs do not reach (Supplementary Fig. S3). *ρ*nc = 0.592±0.011 (pooled 2-seed 5-fold CV, non-conserved genes, gene-operon split with zero cluster leakage). **d,** Monotone deployment ladder: each tier adds a biological input class (protein → + coding DNA → + 5*^′^* untranslated region (UTR) → + full operon → + host genome) and yields a statistically significant *ρ*nc gain (Δ = +0.074 total; Table ED4). The monotone ordering mirrors the biological hierarchy of gene regulation, consistent with multi-scale regulatory structure that the fusion model resolves.

A precomputed biophysical feature vector (69d) bridges these scales with mechanistic signals: codon adaptation indices, translation-initiation energetics, protein physicochemistry, intrinsic disorder, and operon positional features. Under five-fold cross-validation on the gene-operon split with two independent seeds (pooled 10-sample), this ∼25M-parameter model reaches *ρ*_nc_ = 0.592 ± 0.011 on non-conserved genes and *ρ* = 0.667 ± 0.014 overall; split-to-split variance is more than the seed-to-seed variance (around four-fold).On singletons (71% of genes, leak-free by construction), XP5 reaches 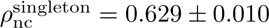 versus ESM-C solo at 0.565 ± 0.012, a Δ = +0.064 fusion advantage independent of any operon-level label-identity confound (Fig. S5).

The operon-level view provides the strongest predictive signal of any single modality (Evo-2 7B solo: *ρ*_nc_ = 0.565). Leave-one-out (LOO) ablation (Extended Data Table ED1; Fig. 3c) shows operon context as the largest contributor (Δ*ρ*_nc_ = −0.031). Genome context ranks third on non-conserved genes (single-seed Δ*ρ*_nc_ = −0.010, multi-partition −0.006; Extended Data Table ED1), despite being negligible by overall *ρ*, suggesting that chromosomal context becomes relevant where gene family-level shortcuts are absent. (Additional recipe variants and ablations in the gene-operon split reiterate these findings, see Supplementary Table S2).

**Fig. 3.**
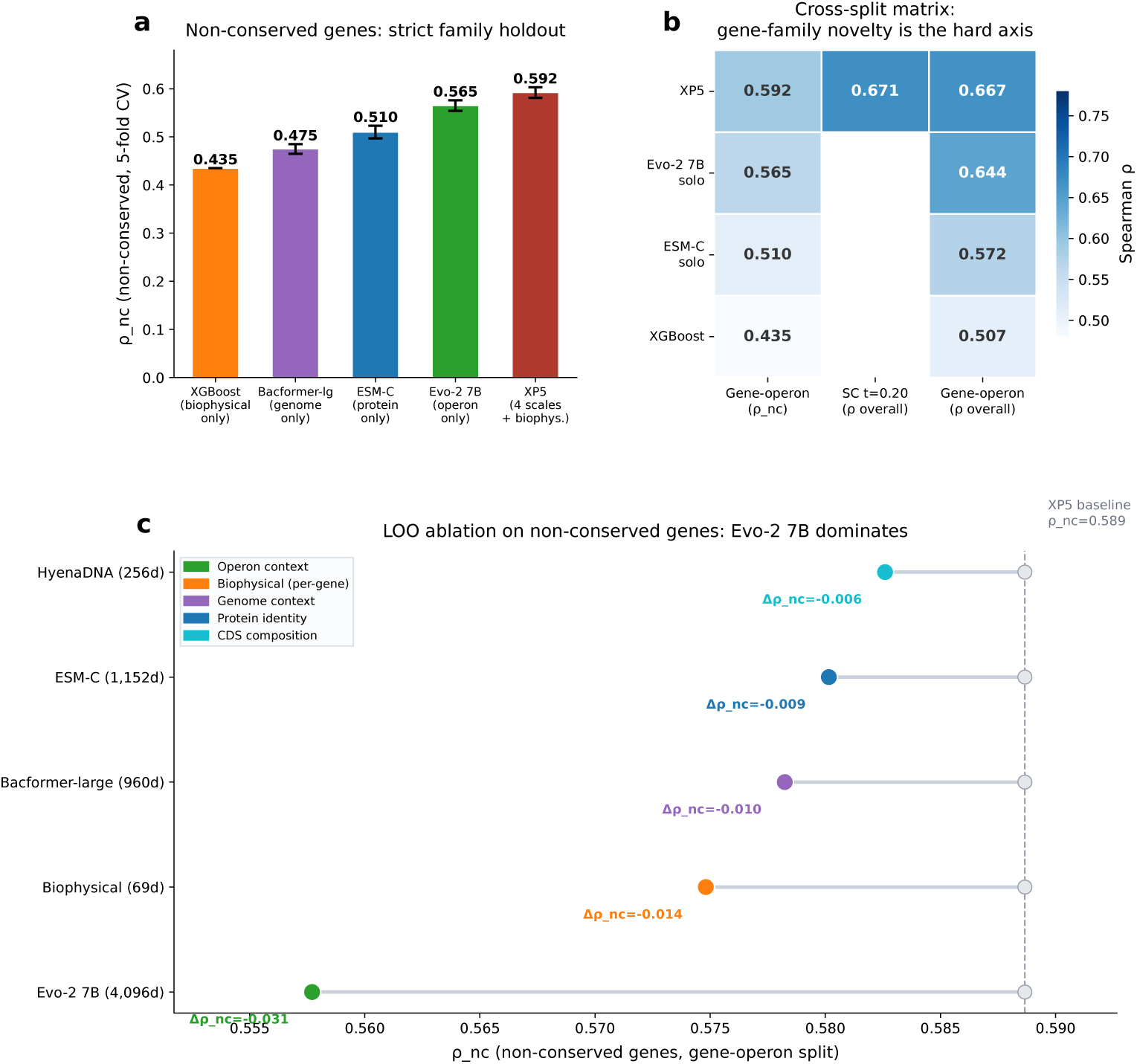
Multimodal fusion improves prediction on non-conserved genes. All panels report *ρ*nc (non-conserved genes) under gene-operon holdout unless noted. **a,** Five levels of model complexity (five-fold CV *ρ*nc): biophysical-only XGBoost (0.435), genome-only Bacformer-large (0.475 ± 0.010), protein-only ESM-C (0.509 ± 0.013), operon-only Evo-2 7B (0.565 ± 0.011), and XP5 (0.592 ± 0.011; Extended Data Table ED3). Each scale alone provides signal; XP5 integrates all four foundation-model scales plus the biophysical block. **b,** Cross-split comparison: *ρ*nc on gene-operon and species-cluster splits, alongside overall *ρ* for reference. **c,** Leave-one-out ablation (Δ*ρ*nc): Evo-2 7B is the major contributor (−0.031), followed by biophysical features (−0.014) and Bacformer-large (−0.010; Extended Data Table ED1).

The ability of a protein language model (PLM) to predict protein expression strictly from the amino acid sequence improves with its inherent size (as measured by number of model parameters); however, this saturated beyond 600M parameters (Fig. 4a). Fusing two same-scale protein models yields a small but statistically significant improvement: ESM-C+ProtT5-XL[35] reaches *ρ*_nc_ = 0.518 (Tier A; +0.009 over ESM-C alone, *p <* 10^−4^); same-scale stacking consistently underperforms cross-scale addition in our experiments, supporting the principle of one foundation model per biological scale. Adding coding-sequence composition (DNA) and biophysical features brings *ρ*_nc_ to 0.530 (Tier B); adding 5*^′^* UTR context to 0.543 (Tier B^+^); full operon DNA to 0.576 (Tier C); and host genome to 0.592 (Tier D). These five tiers form a strictly monotone deployment ladder (Table ED4; Fig. 2d): each step adds a biological input class and yields a significant improvement. Each scale contributes; none alone reaches XP5. The greedy forward-stepwise order by *ρ*_nc_ (operon → genome → CDS → biophysical → protein; Supplementary Fig. S6) is near-inverse to the deployment ladder (protein → CDS → UTR → operon →genome). The two orders are orthogonal: greedy reflects information content, the ladder reflects user-input availability. Rank-averaging five recipes (XP5 plus four near-XP5 variants) adds a further +0.016 on top of XP5 (*ρ*_nc_ = 0.606 ± 0.012, paired *p* = 0.0002; Table ED4 row D_ens_; decomposition in Supplementary Table S4).

**Fig. 4.**
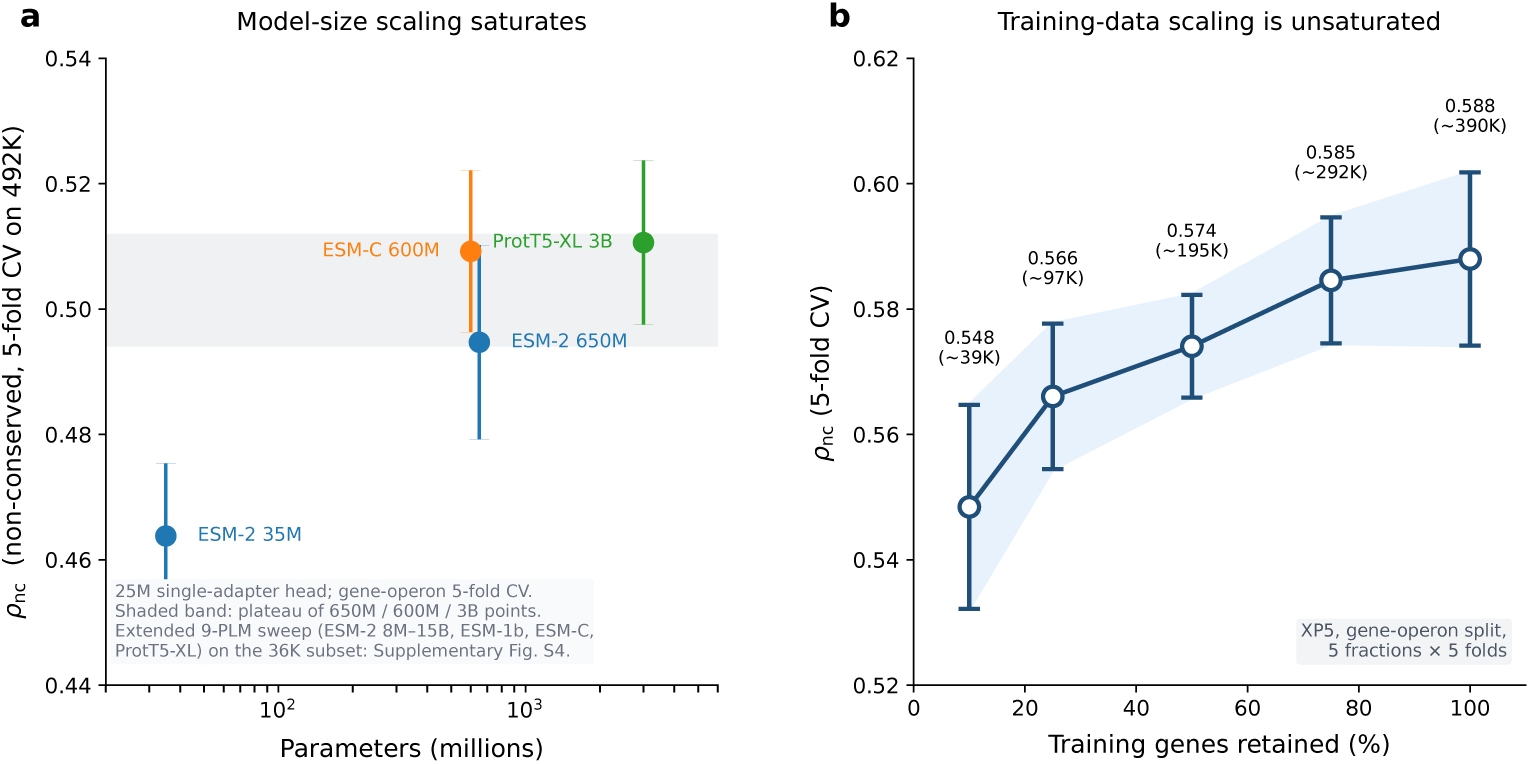
The performance ceiling is data-limited, not model-limited. **a,** Protein-language-model scaling saturates above 600M parameters. Four PLMs (ESM-2 35M, ESM-2 650M, ESM-C 600M, ProtT5-XL 3B) evaluated on the full 492K 5-fold CV with a single-adapter 25M-parameter MLP head (the same head class and capacity as the XP5 production fusion). Error bars = fold s.d. ESM-2 35M (*ρ*nc = 0.464 ± 0.012) anchors the pre-plateau rise; ESM-2 650M (0.495 ± 0.016), ESM-C 600M (0.509 ± 0.013), and ProtT5-XL 3B (0.511 ± 0.013) all sit within one standard deviation of each other across a five-fold parameter spread; these three points define the saturation band. XP5 (*ρ*nc = 0.592, Fig. 3a) sits substantially above this band on the same 492K scale, quantifying the multi-modality fusion gain over any PLM solo at matched head capacity (+0.081 over the best single-PLM point). An extended 9-point ESM-2 family sweep (8M→15B) on the 36K curated subset at the same head scale reproduces the plateau and rules out “starved head” as a confound (Supplementary Fig. S4). **b,** XP5 training-fraction scaling is unsaturated. *ρ*nc rises monotonically from 0.548 at 10% of training genes (∼39K) to 0.588 at 100% (∼390K) across five gene-operon folds (mean ± s.d.). Together, panels a,b indicate that the ceiling is set by training data, not protein-representation capacity.

### Performance saturates with model size but not with training data

Two orthogonal scaling axes clarify what limits the predictive ability of the fusion model (Fig. 4). Under a single-adapter 25M-parameter head matching the production head scale and evaluated on the full 492K 5-fold CV, protein-language-model capacity saturates above 600M parameters: ESM-2 650M reaches *ρ*_nc_ = 0.495 ± 0.016, ESM-C 600M 0.509 ± 0.013, and ProtT5-XL 3B 0.511 ± 0.013, all within one standard deviation of each other, despite a five-fold spread in parameter count (Fig. 4a). ESM-2 35M at 0.464 ± 0.012 marks the pre-plateau anchor. An extended capacity sweep on the 36K subset at the same head scale covers the full ESM-2 family (8M→15B), ESM-1b 650M, ESM-C 600M, and ProtT5-XL 3B; ESM-2 15B does not exceed ESM-2 650M even at capacity-matched evaluation, ruling out “starved head” as a confound (Supplementary Fig. S4).

Training-data volume, in contrast, is unsaturated: the XP5 scales monotonically from *ρ*_nc_ = 0.548 at 10% of training genes to 0.588 at 100% (five-fold CV; Fig. 4b). Broader measurements and additional species remain productive paths for improving baseline cross-species transfer; within-species precision is ultimately constrained by condition-specific regulatory dynamics, not by protein-representation capacity. Given that the ceiling is data-limited, we next dissect which of the five modalities carries the predictive signal and whether that attribution is stable across split designs.

### Operon context dominates under both generalization regimes

The gene-operon and species-cluster holdouts test fundamentally different kinds of generalization (novel gene families versus novel species), and the consistency of the operon contribution across both regimes is evidence that this signal reflects transferable regulatory patterns, not a split-design artifact. The loss in prediction performance when the operon representation is omitted from the input (Δ*ρ*_nc_ = −0.031 under gene-operon holdout) is replicated under species-cluster holdout: Δ*ρ*_nc_ = −0.028 ± 0.005 (23-cluster LOSCO, 23/23 clusters negative, *p <* 10^−17^).Ablation studies consistently rank nucleotide-level signals above amino-acid-level signals in this regime (Extended Data Table ED5).

### Cross-species and cross-regime transfer: where the model generalizes and where it does not

Within the species-cluster split at *t* = 0.20, shared-family genes reach *ρ* = 0.724 whereas novel-family genes reach only *ρ* = 0.492 (Fig. 6). Most of the apparent cross-species signal comes from recognizing known gene families, not from learning transferable regulation (Δ*ρ* = 0.232).

Under the most stringent leave-one-species-cluster-out (LOSCO) holdout, where entire phylogenetic clusters are removed, XP5 retains *ρ*_nc_ = 0.580 ± 0.012 (Fig. 6c; Extended Data Table ED2). Per-species leave-one-species-out (LOSO) reveals wider variation, including one outright failure: *P. aeruginosa* PAO1 drops to *ρ*_nc_ = 0.017, and a strict variant that additionally holds out the species-level Abele record yields *ρ*_nc_ = 0.003 (Methods), so the failure is not explained by cross-source label leakage. We do not claim cross-species transfer to PAO1-style proteomics under LOSO. All external evaluations below use the frozen XP5 and tier-locked recipes from the gene-operon benchmark; the external datasets were never seen during model or tier selection, so the numbers reflect true held-out performance. The frozen XP5 model transfers cleanly to three independent, held-out *E. coli* proteomics technologies (*ρ* = 0.61–0.67 under LOSO with *E. coli* withheld during training) and a 56K-gene temporal holdout (*ρ* = 0.657; Fig. ED1b,c). These results establish utility for native-expression prediction within the training distribution. At the boundaries of that distribution, performance degrades in characterizable ways. On a 24,200-variant subset (∼10%) of the Cambray 2018 synthetic-GFP library[27], a held-out benchmark not used during training or selection, classical biophysical features (*ρ* = 0.434) outperform every foundation-model recipe we tested (*ρ <* 0.17; Fig. ED1d).

Cross-gene regulatory prediction on natural transcripts (our training task) and within-gene sequence optimization of a single GFP coding region (the Cambray task) are complementary problems that rely on different signals. Held-out heterologous-expression datasets (Boël, Price/NESG 2011) show that DNA modalities degrade prediction when native operon context is absent (Fig. ED1b). A held-out test on *Synechococcus elongatus*[23] (*ρ* = 0.147; zero *Synechococcus* in training, with only one other cyanobacterium, *Microcystis aeruginosa*, present) shows prediction degrading with phylogenetic distance. The model ranks conserved housekeeping genes correctly as high-expression but does not distinguish lineage-specific high-abundance proteins from their family background, a known failure mode for foundation models trained on under-represented phyla[3]. On two internal construct sets (*n* = 45 periplasmic nanobodies and *n* = 118 cytosolic binders; Aikium internal data), rank-averaging Aiki-XP Tier C with NetSolP solubility exceeds either alone, indicating deployment-ready complementarity with existing solubility tools (Discussion).

To our knowledge no prior method combines this scale, leakage control, and multimodal breadth: existing approaches are either restricted to single species[20, 33, 37], predict mRNA rather than protein[19], or evaluate under different homology thresholds[22]. Re-evaluation on our gene-operon split shows that single-model approaches reach at most protein-PLM performance: a CaLM[32] codon-level linear probe reaches *ρ*_nc_ = 0.519 (Ridge, 20K-gene training subsample on fold 0), matching ESM-C solo (*ρ*_nc_ = 0.509) and confirming that codon tokenization implicitly captures protein information; TXpredict[19], which predicts mRNA rather than protein, reaches *ρ* = 0.411 (*ρ*_nc_ = 0.323). Both fall ≥0.07 *ρ*_nc_ below XP5.

## Discussion

We set out to test whether multi-scale integration across genome, operon, coding-sequence, and protein views improves bacterial expression prediction in regimes where gene memorization via protein-sequence similarity does not suffice (gene-operon and species-cluster splits). Aiki-XP makes two advances along this axis. First, multimodal fusion improves bacterial expression prediction under strict leakage control: the Δ*ρ* ≈ 0.09 gap between random and gene-operon splits (Extended Data Table ED3) shows that benchmarks without family-level control overestimate generalization[1, 2]. Second, operon-level DNA context is the strongest predictor of within-species relative expression in both generalization regimes, with comparable effect sizes (Fig. 5b). The multi-scale system remains valuable because the smaller contributions aggregate to Δ*ρ*_nc_ = +0.025 over operon-only, driving the deployment ladder (Fig. 2d). Individual LOO effects smaller than ∼0.015 are partition-dependent and are therefore reported only with multi-partition confirmation.

**Fig. 5.**
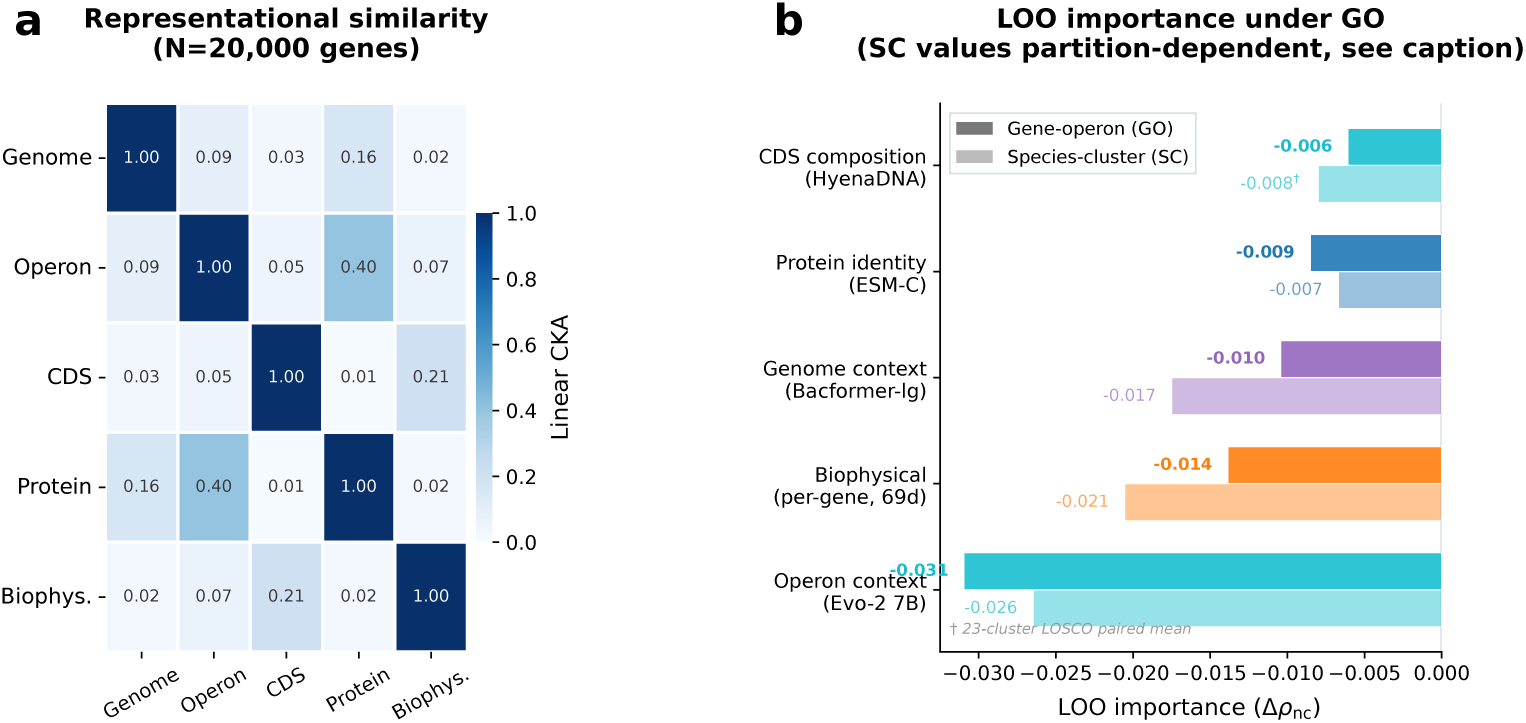
Modality complementarity and LOO importance (XP5). **a,** Linear centered kernel alignment (CKA) between all five XP5 modality groups (*N* = 20,000 genes). The highest inter-scale similarity is between the protein view (ESM-C) and the operon view (Evo-2 7B; CKA= 0.40), reflecting shared protein-coding information encoded through different biological contexts. CDS-level features (HyenaDNA) correlate most with the biophysical block (CKA= 0.21), both capturing codon-level composition. All other inter-scale CKA values are *<* 0.16, confirming genuine complementarity across biological scales. **b,** LOO importance under gene-operon (GO, dark bars; Δ*ρ*nc, 5-fold CV) and species-cluster (SC, light bars; single-seed partition) holdouts. Operon context is the primary contributor under both regimes (GO 5-fold Δ = −0.031; SC 23-cluster LOSCO Δ = −0.028 ± 0.005, 23/23 clusters negative, paired *t*-test *p <* 10^−17^). SC values for the other four scales shown here are from one partition; multi-partition paired tests aggregating 23 LOSCO cluster holdouts with three random SC partitions (*N* = 26) resolve CDS at Δ*ρ*nc = −0.006 ± 0.008 (*p <* 10^−3^) and genome context at −0.006 ± 0.006 (*p <* 10^−4^), correcting the single-partition CDS sign flip (+0.006) and the single-partition genome magnitude (−0.018) shown here. Protein and biophysical SC effects remain partition-dependent. The deployment ladder (Fig. 2d) turns this decomposition into a practical recipe.

**Fig. 6.**
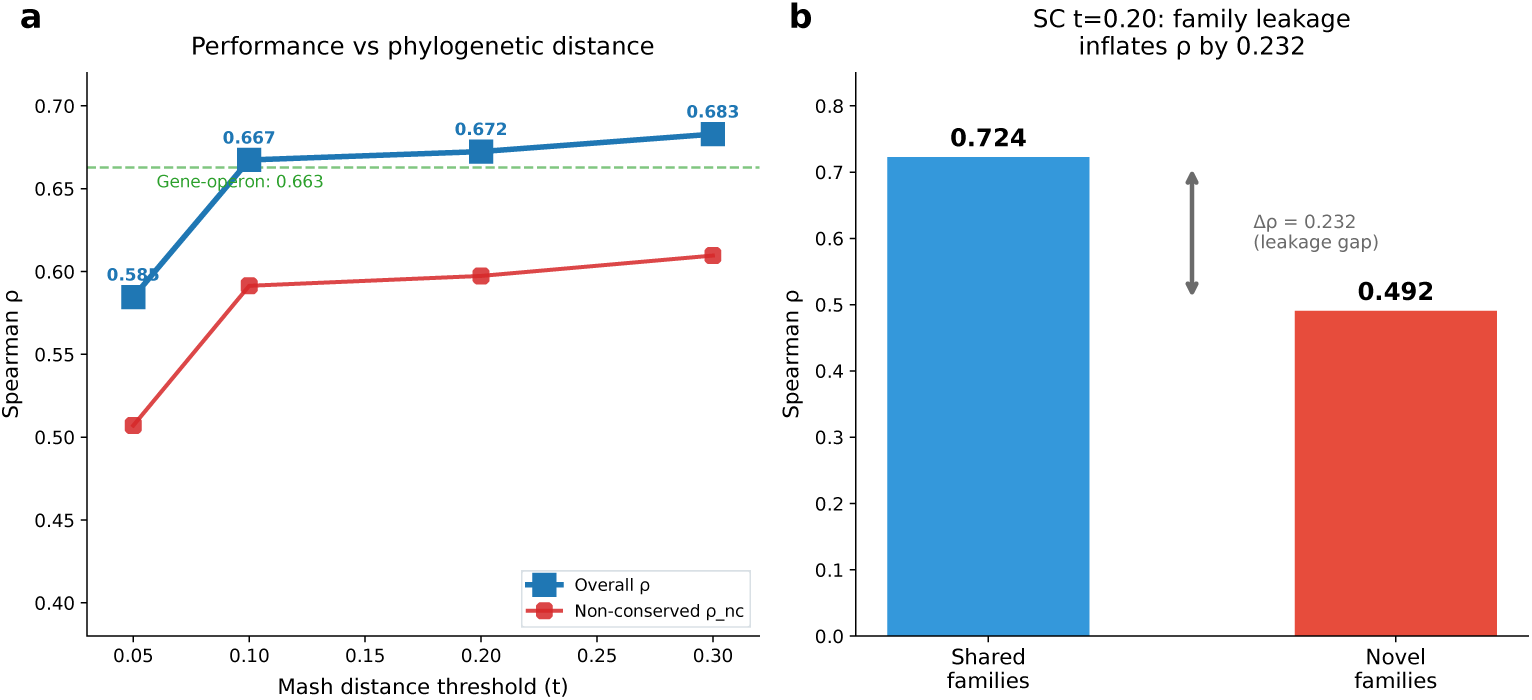
Species-cluster split (XP5): threshold dependence, family recognition, and LOSCO generalization. **a,** Performance under the species-cluster split versus the Mash distance threshold used for single-linkage species clustering and 80/10/10 assignment. Higher *t* yields coarser clusters and more gene families shared across splits. Coarser clusters (higher *t*) share more families across splits; finer clusters (lower *t*) share fewer. At *t*=0.05 (330 fine clusters) test species are phy-logenetically distant from training (*ρ*=0.585); at *t*=0.30 (17 coarse clusters) *ρ*=0.683. Dashed lines: gene-operon reference (the complementary protein-cluster split). **b,** At *t*=0.20, test genes that share a protein-cluster (“family”) with a training gene reach *ρ*=0.724, while genes whose family is absent from the training split reach only 0.492 (Δ*ρ*=0.232). “Gene-operon cluster” and “family” are used interchangeably throughout: both denote the 50%-identity protein cluster used by the gene-operon split. **c,** Leave-one-species-*cluster*-out (LOSCO): 23 phylogenetic clusters held out one at a time; mean *ρ*nc=0.580 ± 0.012. Per-species LOSO across 10 diverse bacteria (Extended Data Table ED2) reveals wider variation than the LOSCO mean suggests: per-species held-out *ρ*nc ranges from 0.434 to 0.683 across nine of ten species (mean 0.566 ± 0.091), with one outright failure (*P. aeruginosa*, *ρ*nc = 0.017; see §Methods).

Protein-level models saturate beyond 600M parameters (Fig. 4a), while operon-level scaling remains productive. On singleton genes, where operon-embedding identity leakage is impossible, XP5 reaches 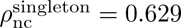 versus ESM-C solo at 0.565 (Δ = +0.064): a leakage-free estimate of the multimodal fusion advantage. Removing the operon view while retaining all other scales recovers no more signal than the operon view alone (*ρ*_nc_ ≈ 0.565); the other scales provide signal only in conjunction with operon context. By design, the operon encoder mean-pools an entire polycistronic unit into one vector, washing out per-gene translation-initiation signals. Per-gene biophysical features (ViennaRNA Δ*G*, Shine–Dalgarno energetics[15]) resolve this gap, explaining why a 69-d block is the second-largest LOO contributor. A separate gap is filled by Bacformer[13], a recent genome-context-aware protein encoder whose chromosomal-neighborhood view contributes the third-largest LOO effect on non-conserved genes (Δ*ρ*_nc_ = −0.010 gene-operon, −0.006 multi-partition species-cluster; Extended Data Table ED1) even with operon DNA, CDS, and protein views all present. This reaffirms that genome-level neighborhood signal is complementary to operon-level transcription-unit signal and the two are not interchangeable. BioLangFusion[29] validated central-dogma fusion on small datasets; at pan-bacterial scale with leakage control, we find that one model per biological scale is sufficient for within-species relative expression.

XP5 has ∼25M parameters and runs in seconds on a laptop GPU (featurization requires foundation models; 16 GB VRAM suffices for all except Evo-2 7B). For recombinant engineering, Aiki-XP is complementary to solubility-focused tools such as NetSolP[21] and MPEPE[20]: on two construct sets (Aikium internal data; *n* = 45 periplasmic nanobodies and *n* = 118 cytosolic binders, scored with the *E. coli* K12 *lacZ* chromosomal context after stripping plasmid-specific promoter elements), rank-averaging Aiki-XP Tier C with NetSolP solubility exceeds either alone (cytosolic: rank-avg *ρ* = 0.505 vs NetSolP-only 0.481 and Tier C-only 0.302; periplasmic: *ρ* = 0.237 vs 0.085 and 0.212), indicating orthogonal signals.

Training with 20% modality dropout preserves peak performance while retaining signal when genome modalities are absent (Extended Data Table ED5); this is the recommended strategy for out-of-distribution tasks or highly engineered constructs where standard biological contexts are disrupted.

Two platform contributions may be of independent interest. The pan-bacterial corpus of 492,026 genes across 385 species with resolved gene↔protein↔genome↔operon mappings and precomputed embeddings from twelve foundation models is released as a standing resource for bacterial expression modeling. The accompanying split scheme addresses a systematic source of error specific to prokaryotic proteomics: because quantification aggregates at the operon or protein-group level, a pure protein-cluster split still admits identical operon embeddings and labels across train and test through co-operonic genes that fall in different protein sequence-based clusters. Our approach is to treat all gene clusters where any member pair shares an operon as an indivisible gene-operon cluster, with this whole cluster of genes being assigned to only one of the data splits. This is how we seal the operon-label identity leakage to meaningfully measure *ρ*_nc_; we offer these data-split annotations as a standard control for any model trained on operon-level expression labels. The 41% conserved ribosomal mega-component (Fig. 1d) makes this silent error concrete: standard benchmarks reward its recognition without measuring regulatory learning. Models trained on genomic or proteomic data should report performance on the non-conserved complement.

Prior fusion work has combined 2–5 modalities at smaller scale[29–31]; we extend this to a sequential input-superset design evaluated under pan-species homology control, with per-step significance testing at every tier. To our knowledge, Aiki-XP is the first system in which a biologically-motivated sequence of input scales adds statistically significant predictive power at every step under strict gene-family homology control at pan-bacterial scale. The design blueprint generalizes to other large-scale multi-modal biological prediction tasks, with domain-informed engineering as the driving principle: per-species z-scoring of labels rather than raw abundance; a gene-operon split that closes the operon-level identity leakage missed by pure protein-cluster splits; explicit handling of conserved mega-components; one foundation model per biological scale, not stacking at the same scale; and singleton vs multi-gene stratification to expose operon-identity leakage. Each is a domain choice, not a hyperparameter-search outcome.

We note the limitations of this work. *Prediction target*: XP5 outputs a per-species z-score rank invariant to linear rescaling across quantification platforms, not absolute copies per cell; cross-species absolute comparisons (“does gene X express more in *E. coli* or *B. subtilis*?”) require per-species calibration and are addressed in follow-up work. *Label heterogeneity*: cross-source agreement is high but per-source biases may remain. *Length confound*: a length-only baseline reaches |*ρ*| = 0.067 (∼10% of XP5’s magnitude). *Positional information loss*: all embeddings are mean-pooled; per-residue features[29] could capture local regulatory motifs. *Heterologous expression*: on Boël and Price/NESG, solubility-focused tools (NetSolP, MPEPE) outperform Aiki-XP (Extended Data Table ED4), as expected given these benchmarks measure folding and solubility rather than native transcription/translation. Internal recombinant tests show Aiki-XP correlates more strongly with expression for periplasmic than cytosolic constructs, and does not predict the soluble-vs-inclusion-body outcome (a folding/quality-control endpoint absent from our training signal). Despite these constraints, the results support modeling bacterial gene expression as a multi-scale regulatory process rather than a single-sequence prediction task.

Three directions now look most promising for raising the ceiling. First, the training-data axis is unsaturated (Fig. 4b), so broader proteomics coverage across growth conditions and under-represented phyla should yield further gains. Second, condition-aware extensions and per-species absolute-abundance calibration open a path to downstream applications in real-world protein and genetic engineering.

Third, with larger, better-annotated datasets, regulatory interactions should become more patent, justifying scaled-up Bacformer-like models that absorb genome-level information to complement the operon-level context that likely remains the strongest independent predictor.

## Online Methods

### Dataset construction

Expression labels from PaXDb v6.0[4] (multi-study integrated proteomics; ppm units) and Abele et al. 2025[5] (single-study data-dependent acquisition (DDA) iBAQ; Orbitrap Exploris 480) were per-species z-scored and combined into a frozen production table of 492,026 genes across 385 species. Per-species z-scoring is necessary because raw abundance values span *>*5 orders of magnitude and differ systematically between species and quantification technologies (Supplementary Fig. S2); all *ρ* values in this paper are computed on z-scored predictions and labels. Within each species, Spearman *ρ* is exactly invariant to z-scoring (it is a rank statistic), and Pearson *r* is also invariant because z-scoring is an affine transform; the cross-species *ρ* on raw-reconstructed predictions is higher (0.715 versus 0.648 on the same Abele test rows where a raw log_2_ iBAQ scale is defined; Δ*ρ* = +0.067) because it includes the species-mean component, which is itself a measurement artifact: per-species mean abundance does not correlate across independent platforms (*ρ* = 0.23, *p* = 0.25; 27 species measured by both PaXDb and Abele; Supplementary Fig. S2). Direct cross-source label agreement on 25,844 genes from 27 species covered by both sources is *ρ* = 0.932; z-scored labels are highly concordant despite different quantification pipelines; this agreement is identical whether computed on z-scored or raw values within each species. Label-treatment robustness check (fold 0 of the gene-operon split, single seed): replacing per-species z-score with a per-species rank-to-standard-normal transform yields pooled *ρ*_nc_ = 0.586 versus the XP5 baseline 0.589, within seed-level noise. Source-restricted training (XP5 trained on PaXDb genes alone or on Abele genes alone; fold 0, single seed) recovers per-source pooled *ρ*_nc_ comparable to the combined-source XP5 when each model is evaluated on its own source’s test rows (PaXDb test: 0.619 vs 0.612; Abele test: 0.567 vs 0.564); the value of the 492K-gene combined corpus is taxonomic coverage, not per-species accuracy. A within-species label-shuffling test shows that 99% of XP5’s *ρ* = 0.663 is genuine gene-level prediction, not between-species structure (Supplementary Fig. S8).

### Feature modalities

We extracted thirty feature modalities (∼24,100 total dimensions) across twelve neural models, from which XP5 retains one per biological scale (see Results). Extracted models: ESM-2 650M (1,280d)[7], ESM-C 600M (1,152d), ProtT5-XL 3B (1,024d)[35], HyenaDNA (256d)[8], DNABERT-2 (768d)[9], Nucleotide Transformer v2 500M (1,024d)[10], CodonFM 1B (2,048d), RiNALMo (1,280d)[11], 5*^′^*UTRBERT (768d), Bacformer-base (480d) and Bacformer-large (960d)[13], and Evo-2[12] (1B: 5,120d; 7B: PCA-4096 from 11,264d full-operon embeddings). A precomputed biophysical feature vector (69d total) captures mechanistic signals that foundation models do not learn from sequence alone: host-specific codon adaptation indices (tAI, CAI, ENC; 11d)[14], translation-initiation energetics from each gene’s 5*^′^* window (mRNA folding free energy and Shine–Dalgarno binding Δ*G* via ViennaRNA[15]; SD–AUG spacing penalty[16]; 16d)[17, 34], protein physicochemistry (pI, hydropathy, instability index; 24d), intrinsic disorder and charge patterning (8d), and operon positional architecture (gene position, intergenic distances; 10d). The translation-initiation and operon-position features are computed per gene and provide gene-specific signals that the operon-level mean-pooled embedding cannot resolve. This feature vector enters the greedy selection as a single block. XP5 uses Evo-2 7B full-operon PCA-4096 in the DNA slot. Modality coverage is ≥99.8% across all genes; residual gaps were zero-filled during featurization.

### Split construction

*Gene-operon split*: MMseqs2[18] at 50% identity yields 129,078 protein clusters, none of which span train/val/test by construction. Clusters are then merged by Union-Find over a bipartite graph with two edge types — shared compound operon identity and shared protein sequence. Every non-conserved component (*<*5,000 genes; 59% of the dataset) is assigned to a single split as an indivisible unit. No non-conserved operon therefore straddles the split, so operon-level label-identity leaks cannot enter *ρ*_nc_. The single mega-component (the 41% conserved ribosomal cluster, Fig. 1d) is split at cluster granularity under an operon-coherence penalty; any residual operon fragmentation is confined to this component and excluded from *ρ*_nc_ by definition. Train/val/test: 390,640/50,694/50,692 genes. (scripts/protex/build_hard_production_split.py). *Species-cluster split*: Pairwise Mash distances were computed using sourmash (k=31, scaled=1000), and species were grouped by single-linkage clustering at thresholds 0.05–0.30. At *t* = 0.20, the split contains 300 training, 37 validation, and 48 test species. *Random split*: species-proportional stratified random assignment at the gene level.

### Conserved-component annotation

Union-Find on a bipartite graph linking protein clusters to compound operon identifiers identified one 202,945-gene component (41.2%) spanning all species.

### Modeling

Single-adapter architecture: concatenated mean-pooled embeddings → per-modality pyramid adapters → fusion MLP → scalar output. Parameter budget: ∼25M (capacity-matched). AdamW, lr 2 × 10^−4^, cosine schedule, 200 warmup steps, batch size 64, patience-10 early stopping on validation Spearman *ρ*. A systematic sweep of capacity (10–100M trainable parameters), alternative fusion architectures (seven types), cross-attention, noise-robust losses (six methods), and adversarial domain-adaptation curricula did not exceed the XP5 limit at *ρ* ≈ 0.63 overall (Supplementary Table S2); the *ρ* gain came from scaling the DNA foundation model (1B→7B).

### Metrics

Spearman *ρ* at four strata: overall, non-conserved, cluster-weighted (≥5 test genes per cluster), and per-species (≥10 test genes). Species-cluster splits additionally partition test genes into shared-family and novel-family subsets. Error bars throughout the text are the five-fold standard deviation (ddof=1) unless stated otherwise; 95% confidence intervals on pooled-5-fold estimates (Extended Data Table ED4 footnote) are computed by 2000-resample paired bootstrap on the concatenated test predictions (n ≈ 145,000 per recipe), equivalent to Spearman via rank-then-Pearson. Seed stability is reported by retraining XP5 and tier-lock recipes at a second random seed on the same five gene-operon folds; the seed-level variance is approximately four times smaller than the fold-level variance, so the reported ±fold error bars conservatively reflect the dominant source of variability.Paired *t*-tests across the five folds are reported alongside Wilcoxon tests for robustness, using the standard SciPy implementation. Per-species *ρ* values reported in this paper (Supplementary Figs. S2b, S8b) are computed by pooling each species’ predictions across all five gene-operon CV folds and computing one *ρ* per species (Method 2); the alternative of computing a per-species *ρ* within each fold and then aggregating across folds (Method 1) yields a median per-species cross-fold standard deviation of ∼0.10 driven by small per-fold sample sizes, an order of magnitude larger than the headline pooled-*ρ* five-fold s.d. of ∼0.012, and is reserved for cross-fold stability diagnostics rather than headline per-species reporting. Per-gene regression calibration of the XP5 (20 quantile bins of predicted *z*-score across five gene-operon test folds) gives mean reliability MAE 0.069 ± 0.030 and slope 0.94 ± 0.03, consistent with mild under-confidence at distribution tails; regression MSE is 0.52 ± 0.02 overall and 0.57 ± 0.02 on non-conserved genes.For external prediction on a species not represented in training, raw model outputs are uncalibrated *z*-scores against the implicit training-distribution mean (per-species *µ̂_S_* and *σ̂_S_* are not stored at inference time); when ground-truth abundances are available for a panel of measured proteins from the target species, an affine recalibration (*y*_cal_ = *a y*_raw_ + *b*, fit by ordinary least squares on the panel) corrects the per-species offset and slope (scripts/protex/xp5_calibration_kit.py).

Spearman *ρ* is invariant to this affine transform, so all rank-based metrics in this paper are unaffected; the recipe affects only absolute-magnitude predictions.

### Per-species LOSO failure mode (*P. aeruginosa*)

The single notable LOSO failure (Extended Data Table ED2) is for *P. aeruginosa*. In our combined dataset *P. aeruginosa* is represented by two independent label records from different upstream proteomics pipelines: 5,169 PAO1 genes (taxid 208964) drawn from PaXDb v6.0[4] integrated with supplementary single-source measurements from GPMDB and MassIVE, and 1,846 *P. aeruginosa* species-level genes (taxid 287) from the Abele[5] single-study iBAQ atlas. The default LOSO setup holds out the PAO1 record only and yields *ρ*_nc_ = 0.017 on the held-out PAO1 genes; a strict variant that additionally holds out the species-level Abele record yields *ρ*_nc_ = 0.003. Removing the cross-source overlap therefore does not improve the failure, indicating that for this species the limit is not label leakage between the two records. Independent validation on the V2.1 temporal holdout reproduces the failure pattern at PAO1 (*ρ*_ensemble_ = 0.277, ranking 4 of 89 species evaluated), while other *Pseudomonas* taxa rank near the median. We do not claim XP5 generalizes to PAO1-style proteomics under either LOSO regime.

### Computational resources

The full experimental campaign comprises ∼1,600 training runs. Fusion model training ran on four NVIDIA RTX 5000 Ada Generation laptop GPUs (16 GB VRAM); each run took ∼15 minutes (∼350 GPU-hours total). Foundation-model embedding extraction used three NVIDIA A100 SXM4 nodes (two 40 GB and one 80 GB, on Google Cloud Platform), with Evo-2 7B extraction (∼62 h on an A100-80GB) the single largest cost. Inference with the frozen XP5 takes *<*5 s per proteome on a laptop GPU. Estimated total carbon footprint: ∼38 kg CO_2_e (ML CO_2_ Impact calculator, 0.11 kg CO_2_/GPU-hr for cloud; Google Cloud Platform us-central1 carbon intensity).

### Use of AI tools

Claude (Anthropic) and ChatGPT (OpenAI) were used as coding assistants for data pipeline development, embedding extraction scripts, statistical analysis, and manuscript editing. All AI-generated code was reviewed, tested, and validated by the authors. All quantitative claims were independently verified against source result files in the code repository. Scientific hypotheses, result interpretation, and experimental design were the sole responsibility of the authors.

## Data availability

The training dataset (492,026 genes × 24 columns), gene-operon and species-cluster split files, pre-extracted foundation-model embeddings for all twelve evaluated models on the 492K set and external benchmarks, and a manifest of the 1,831 reference genomes used are deposited at Zenodo together with all trained checkpoints and evaluation JSONs under a single combined DOI: 10.5281/zenodo.19639621 (shared with the Code availability deposit below). Expression data were obtained from PaXDb v6.0[4] (https://pax-db.org) and the Abele proteomics atlas[5] (ProteomicsDB Project 4498). Supplementary single-source measurements incorporated into the training set: *Cronobacter sakazakii* and *Shewanella putrefaciens* from MassIVE MSV000096603 (NSAF-normalized iBAQ values derived from the raw MaxQuant output of the same Abele lab study, fewer species than the main atlas); *Legionella pneumophila* from GPMDB accession 272624 (spectral counts, August 2014 snapshot). All supplementary sources were per-species z-scored into the same normalized scale as PaXDb. External benchmarks (none used during XP5 training or tier selection; all held out until final evaluation): Boël 2016 (Northeast Structural Genomics Consortium (NESG)/TargetTrack), Price/NESG 2011 (PMID 21455294), Seattle Structural Genomics Center for Infectious Disease (SSGCID; TargetTrack), *Synechococcus elongatus* data-independent acquisition mass spectrometry (DIA-MS)[23] (PRIDE PXD062851), Cambray 2018 GFP library[27] (Supplementary Data 15), and three held-out *E. coli* K12 proteomics technologies used for the LOSO tier comparison (Fig. ED1b): Li 2014 ribosome profiling[24], Mori 2021 DIA-MS[25], and Taniguchi 2010 single-cell YFP[26]. Full provenance for every data source is documented in DATA_SOURCES.md within the Zenodo deposit.

## Code availability

Source code for data assembly, split construction, feature extraction, model training, evaluation, and figure reproduction is available at https://github.com/aikium-public/aiki-xp and archived at Zenodo under the same combined DOI cited in Data availability (10.5281/zenodo.19639621). Trained model checkpoints (five-fold ensemble for all deployment tiers), PCA transforms, and result JSON files backing every quantitative claim are included in the same Zenodo deposit. The repository also provides two Docker images: an inference-only image that scores pre-extracted embeddings, and an end-to-end image that extracts ESM-C and ProtT5-XL embeddings from raw FASTA input and runs Tier A prediction on CPU. For inference on query proteins with engineered tags (His-tags, HiBit, FLAG, myc, linker sequences, or non-native signal peptides), the canonical normalization routine sequence_normalization.py strips tags before featurization. For recombinant *E. coli* constructs expressed from lac-regulated vectors, Tier C inference can be performed by wrapping the coding sequence in the upstream 100 nt and downstream 50 nt of the chromosomal lacZ locus in K12 MG1655; documentation for this protocol is included in the repository.

## Acknowledgements

We thank the PaXDb and Abele teams for making their proteomics data publicly available, and the curators of MassIVE, GPMDB, ProteomeXchange, NCBI GenBank and the Entrez system, DOOR2, RegulonDB, and MicrobesOnline for maintaining the open bioinformatics infrastructure that made this cross-database integration possible. We thank the developers of all foundation models evaluated in this work: ESM-2 and ESM-C (Meta), Evo-2 (Arc Institute), ProtT5-XL (TUM/Rostlab), DNABERT-2, HyenaDNA, Bacformer, CodonFM, Nucleotide Transformer v2, RiNALMo, and 5*^′^*UTRBERT, for open model weights; the ViennaRNA team for the RNA folding toolkit; the authors of MMseqs2 and Mash/sourmash for the sequence-clustering and genome-distance tools on which our homology-controlled splits depend; and the broader biophysical-features community for releasing open software. A portion of this work was enabled by Google Cloud Platform credits generously provided by the Google for AI Startups Cloud Program.

## Author contributions

H.T. curated the initial version of the dataset, implemented and evaluated the first version of the multimodal fusion model, and provided the basis for the manuscript. R.S.M. contributed to the problem formulation and modeling discussions, provided critical methodological feedback, and helped rewrite the manuscript. S.S. was involved in the conceptual stage of the project, provided the domain knowledge for the correct interpretation of data and results, and helped rewrite the manuscript. V.M. conceived the project, expanded the dataset, implemented the final augmented version of the model, performed the experiments, compiled the results and edited the final manuscript.

## Extended Data

**Table ED1.**
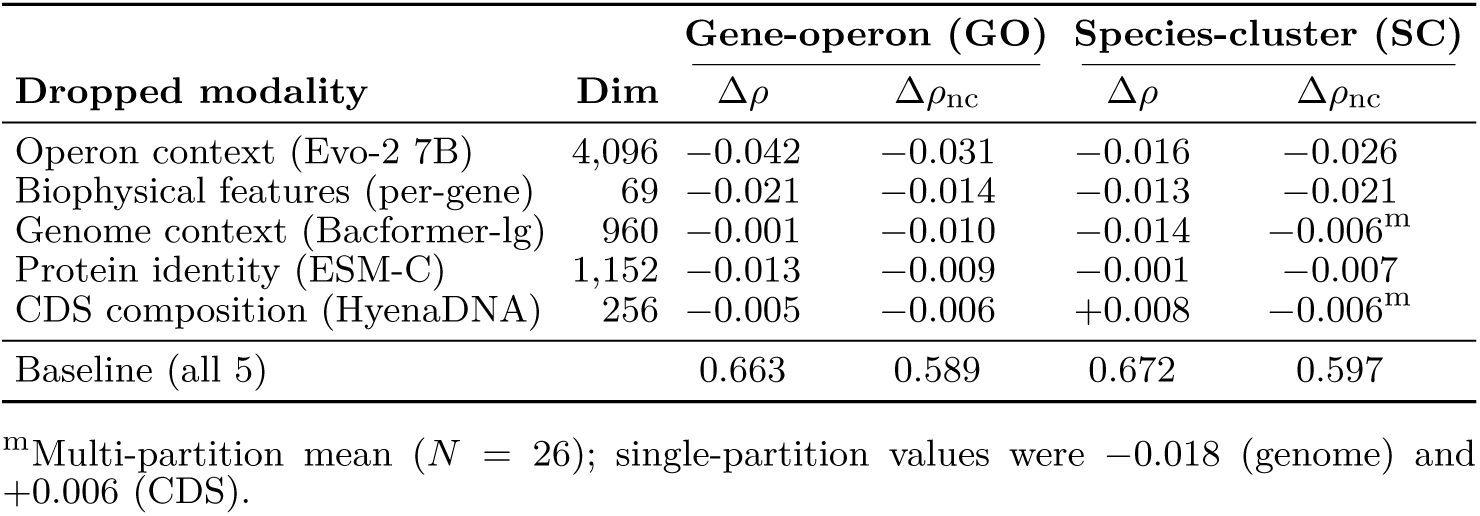
LOO ablation across evaluation regimes. Each row shows the performance drop when one modality is removed, under gene-operon (GO, 5-fold CV) and species-cluster (SC) holdouts. GO values are single-seed (seed 42) so the relative magnitudes are directly comparable across rows; the matched single-seed XP5 baseline is *ρ* = 0.663, *ρ*nc = 0.589. The pooled 2-seed XP5 reported in the abstract and Table ED3 (*ρ* = 0.667, *ρ*_nc_ = 0.592) is the same model evaluated with one additional seed. SC Δ*ρ*_nc_ values for CDS and genome context are multi-partition means (*N* = 26: 23 LOSCO cluster holdouts + 3 random SC partitions); SC values for operon, protein, and biophysical are from a single-seed partition (effects below ≈0.015 are partition-dependent).

| Dropped modality | Dim | Gene-operon (GO) |  | Species-cluster (SC) |  |
| --- | --- | --- | --- | --- | --- |
| | | $\Delta\rho$ | $\Delta\rho_{nc}$ | $\Delta\rho$ | $\Delta\rho_{nc}$ |
| Operon context (Evo-2 7B) | 4,096 | -0.042 | -0.031 | -0.016 | -0.026 |
| Biophysical features (per-gene) | 69 | -0.021 | -0.014 | -0.013 | -0.021 |
| Genome context (Bacformer-lg) | 960 | -0.001 | -0.010 | -0.014 | -0.006<sup>m</sup> |
| Protein identity (ESM-C) | 1,152 | -0.013 | -0.009 | -0.001 | -0.007 |
| CDS composition (HyenaDNA) | 256 | -0.005 | -0.006 | +0.008 | -0.006<sup>m</sup> |
| Baseline (all 5) |  | 0.663 | 0.589 | 0.672 | 0.597 |

<sup>m</sup>Multi-partition mean ( $N = 26$ ); single-partition values were -0.018 (genome) and +0.006 (CDS).

**Fig. ED1.**
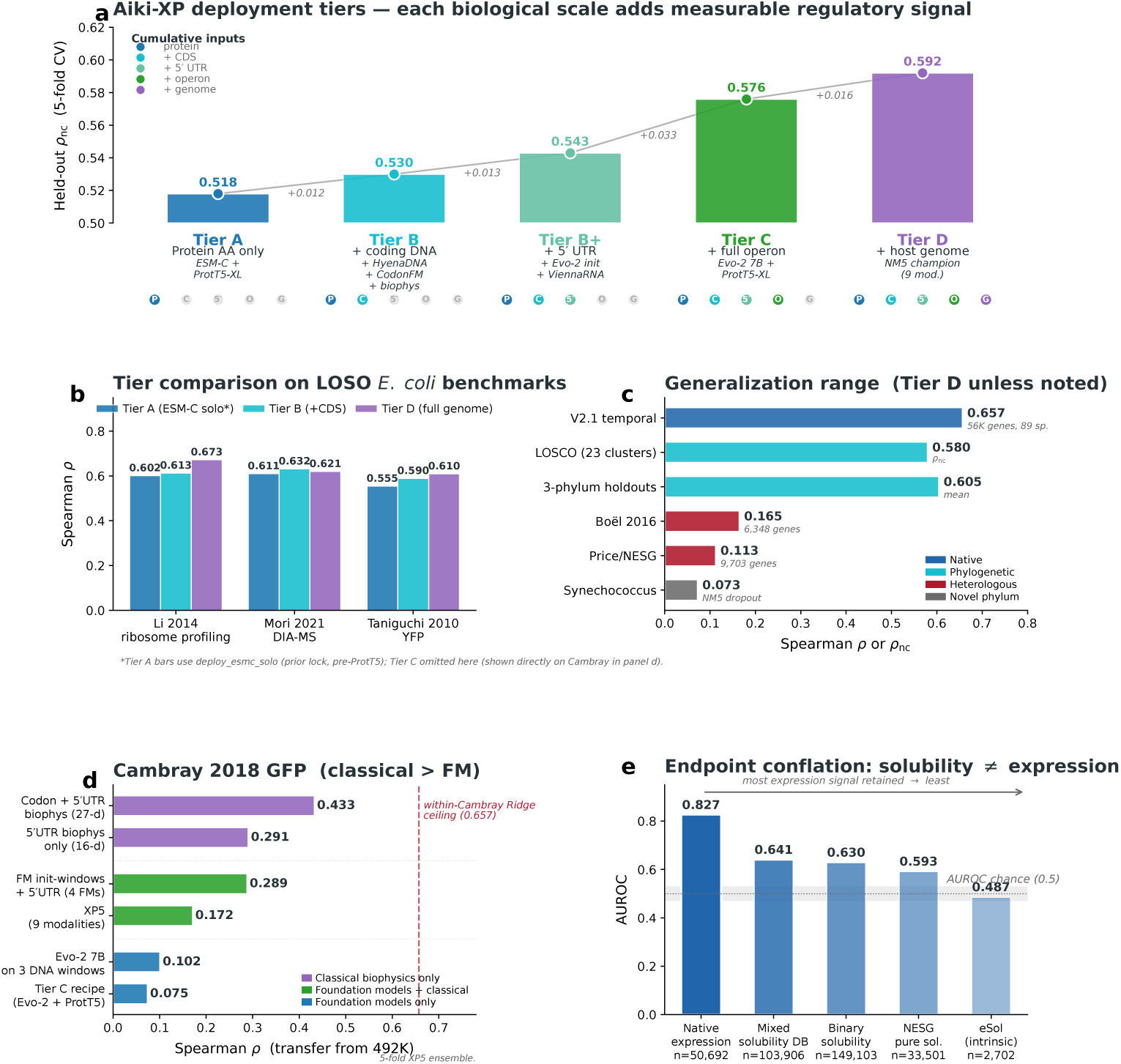
Deployment tiers, native-expression validation, generalization limits, Cambray dilution, and solubility endpoint conflation. XP5 and every tier recipe shown here were locked on the gene-operon benchmark before external evaluation; the datasets in panels **b**–**e** were never used for training or model selection, so every number reports true hold-out performance. **a,** Five Aiki-XP deployment tiers (A → B → B^+^ → C → D) match the model to the user’s available data along a strictly monotone ladder from protein sequence alone to full native genome; each tier’s input is a strict superset of the previous one, and every step adds a statistically significant Δ*ρ*nc. **b,** Tier-stratified comparison on three held-out *E. coli* proteomics datasets under LOSO (Li 2014 ribosome profiling[24], Mori 2021 DIA-MS[25], Taniguchi 2010 single-cell YFP[26]; all *E. coli* K12 genes withheld during training): each additional context type yields a monotone improvement, with Tier D reaching *ρ* = 0.61–0.67 and Tier A *ρ* = 0.56–0.61. **c,** Tier D generalizes across the V2.1 temporal holdout (*ρ* = 0.657, 56K genes added to PaXDb and Abele after the training freeze), 23-cluster LOSCO (*ρ*nc = 0.580), three-phylum holdouts (*ρ* ≈ 0.605) and a held-out *Synechococcus elongatus* test (*ρ* = 0.147; 68% protein-level homology with the training set via conserved gene families, Supplementary Table S3), and retains partial signal on heterologous recombinant expression (Boël T7/BL21 *ρ* = 0.165, Price/NESG cell-free *ρ* = 0.113, both with the Tier A recipe). **d,** Cambray 2018 case study: *N* = 24,200 variants, a ∼10% subset of the full 244,000-variant synthetic GFP library. All bars are 5-fold XP5Ensemble transfers of recipes trained only on the 492K training set; no recipe is trained on Cambray labels. A 27-d classical vector of codon composition plus ViennaRNA 5*^′^* UTR thermodynamics reaches *ρ* = 0.434, higher than every foundation-model recipe we tested, including a 9-modality kitchen-sink variant (*ρ* = 0.172) and pure-FM transfers (*ρ* = 0.07–0.10). Dashed red line: the best within-Cambray ridge fit on the same constructs (*ρ* = 0.657, Evo-2 7B 70-nt initiation-window features), quantifying the upper bound achievable when the Cambray labels are used for training rather than only for evaluation. Full dilution-matrix discussion in Results. **e,** Endpoint gradient. Bars are ordered left to right from the model’s native task to the endpoint that retains least native-expression signal: (i) native bacterial expression (XP5 predictions on the gene-operon 5-fold hold-out, *N* = 50,692; a binary target is formed by dichotomizing the z-scored ground-truth label at its median, and AUROC is computed for the model’s continuous predictions against this high-vs-low binary label, yielding AUROC = 0.827); (ii) the mixed solubility hybrid DB (eSOL + NESG + TargetTrack, AUROC = 0.641); (iii) a binary-solubility aggregate (aggregated public recombinant datasets; AUROC = 0.630); (iv) pure recombinant solubility from the NESG structural-genomics consortium (AUROC = 0.593); and (v) the small *E. coli* intrinsic-solubility set eSol (AUROC = 0.487). Bar opacity tracks the same gradient. The model retains substantial signal across protocols that mix expression with solubility, and only collapses to chance on eSol, the expected pattern for a model trained on native proteomics abundance.

**Table ED2.**
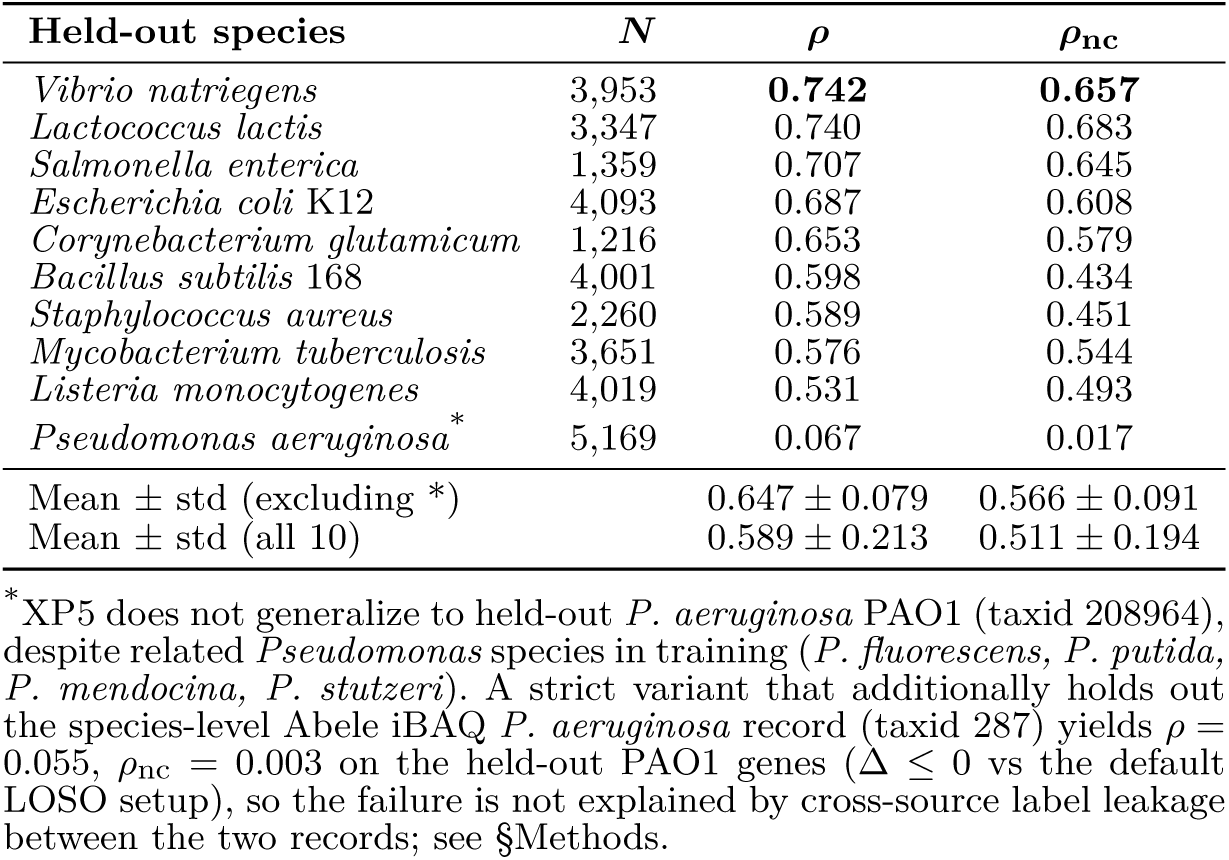
Leave-one-species-out holdout (XP5), per-species held-out. *ρ*. *ρ* and *ρ*_nc_ are computed on the held-out species’ genes only (*N* shown), not on the full LOSO-experiment test set (which is dominated by non-held-out species). Earlier versions of this table inadvertently reported the full-test pooled values; the corrected per-species view exposes wider variation and one outright failure mode.

**Table ED3.** Recipe comparison across the three split regimes. Five-fold CV on the gene-operon split with zero protein leakage; the two right-hand columns show the XP5 recipe evaluated on the species-cluster (*t*=0.20) and random splits for the same data (3 seeds each). The Δ*ρ* ≈ 0.09 gap between random and gene-operon splits is the direct evidence for family-level leakage control: random-split benchmarks overstate generalization by almost 0.1 *ρ*.

| Config | $\rho_{nc}$ (5-fold) | $\rho$ (5-fold) | SC $t=0.20$ ($\rho$) | Random ($\rho$) |
| --- | --- | --- | --- | --- |
| **XP5** | **0.592 ± 0.011** | **0.667 ± 0.014** | 0.671 ± 0.008 | **0.754 ± 0.001** |
| F10 (legacy 10-input baseline)<sup>g</sup> | 0.588 ± 0.011 | 0.664 ± 0.012 | — | — |
| BF-large swap (4 mods) | 0.586 ± 0.012 | 0.662 ± 0.011 | — | — |
| Evo-2 7B+ProtT5 (Tier C) | 0.576 ± 0.011 | 0.648 ± 0.012 | — | — |
| Evo-2 7B solo | 0.565 ± 0.011 | 0.644 ± 0.014 | — | — |
| ESM-C+ProtT5 (Tier A) | 0.518 ± 0.011 | 0.579 ± 0.011 | — | — |
| ESM-C solo | 0.509 | 0.572 | — | — |
| XGBoost baseline<sup>h</sup> | 0.435 | 0.507 | — | — |

<sup>g</sup>F10 is the pre-XP5 champion (10 feature inputs via forward-stepwise selection with Evo-2 1B in the CDS slot); retained here for historical comparison. <sup>h</sup>XGBoost on the 69-d biophysical feature block (single-run fold-0 evaluation, $n = 50,692$ test genes; no 5-fold CV).

**Table ED4.**
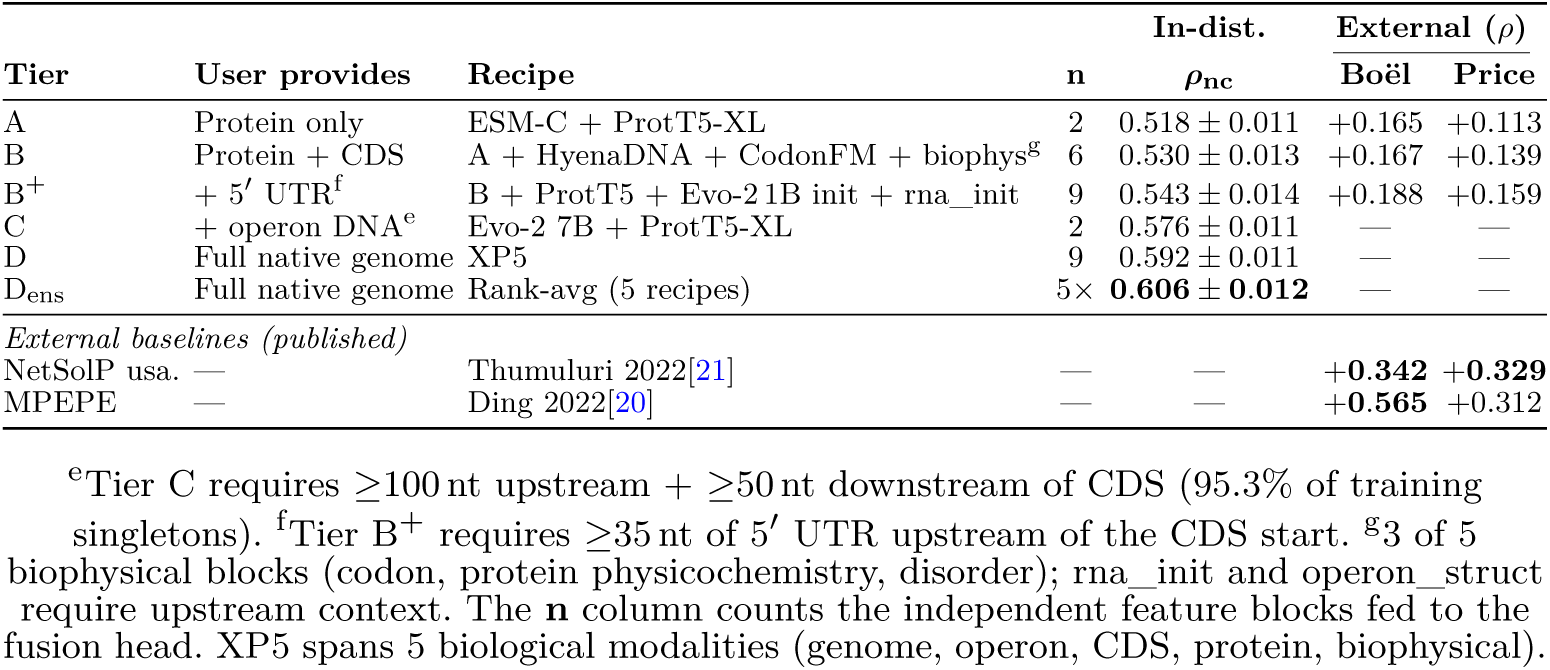
Deployment tiers: in-distribution 5-fold CV *ρ*_nc_ alongside external transfer *ρ* on heterologous recombinant benchmarks. Each tier’s Recipe column lists its locked modality composition. In-distribution selection follows the simplest-not-statistically-worse rule (paired *t*-test, *p <* 0.05 across 5 folds) applied over candidate recipes at each tier. Under this rule, ProtT5-XL upgrades Tier A (Δ= + 0.008, pooled 2-seed *p <* 10^−4^) and simplifies Tier C (4→2 modalities, *p* = 0.124, not significant). Tier B^+^ stacks ProtT5-XL, an Evo-2 1B standalone init-window embedding, and a 16-d ViennaRNA 5*^′^* UTR biophysical block on top of Tier B (Δ = +0.012, *p* = 0.007, 5/5 folds); the three additions are synergistic; none contributes significantly alone. Tiers B and D are unchanged (all ProtT5 additions not statistically significant at those tiers). The Tier D_ensemble_ row is a rank-average of three XP5 retrains plus two fusion variants (Δ = +0.016 vs single-seed XP5, *p* = 0.0002; Supplementary Table S4). The rightmost two columns show the same tier recipes transferred to Boël 2016 (*n*=6,348) and Price/NESG 2011 (*n*=9,703) heterologous recombinant-expression benchmarks via xp5_inference.py, head-to-head against the published NetSolP usability and MPEPE codonC4 baselines. Tiers C and D are not evaluated externally: both require native-genome operon or chromosomal context that recombinant constructs lack.

**Table ED5.** Missing-modality robustness and label noise (XP5). Single-seed (seed 42); the matched-seed all-five-modalities baseline is *ρ* = 0.663, *ρ*nc = 0.589.

| Missing-modality condition | $\rho$ | $\rho_{nc}$ |
| --- | --- | --- |
| All 5 modalities | 0.663 | 0.589 |
| No biophysical | 0.642 | 0.575 |
| DNA only | 0.642 | 0.571 |
| Protein + DNA | 0.647 | 0.578 |
| Protein only (ESM-C) | 0.570 | 0.509 |
| Biophysical only | 0.507 | 0.435 |
| **Label noise (XP5)** |  |  |
| 10% corruption | 0.660 | 0.582 |
| 50% corruption | 0.620 | 0.552 |
| 100% (scrambled) | 0.079 | 0.062 |
| Noise: random permutation of $k\%$ of training labels. 100% is the negative control. | | |

**Fig. ED2.**
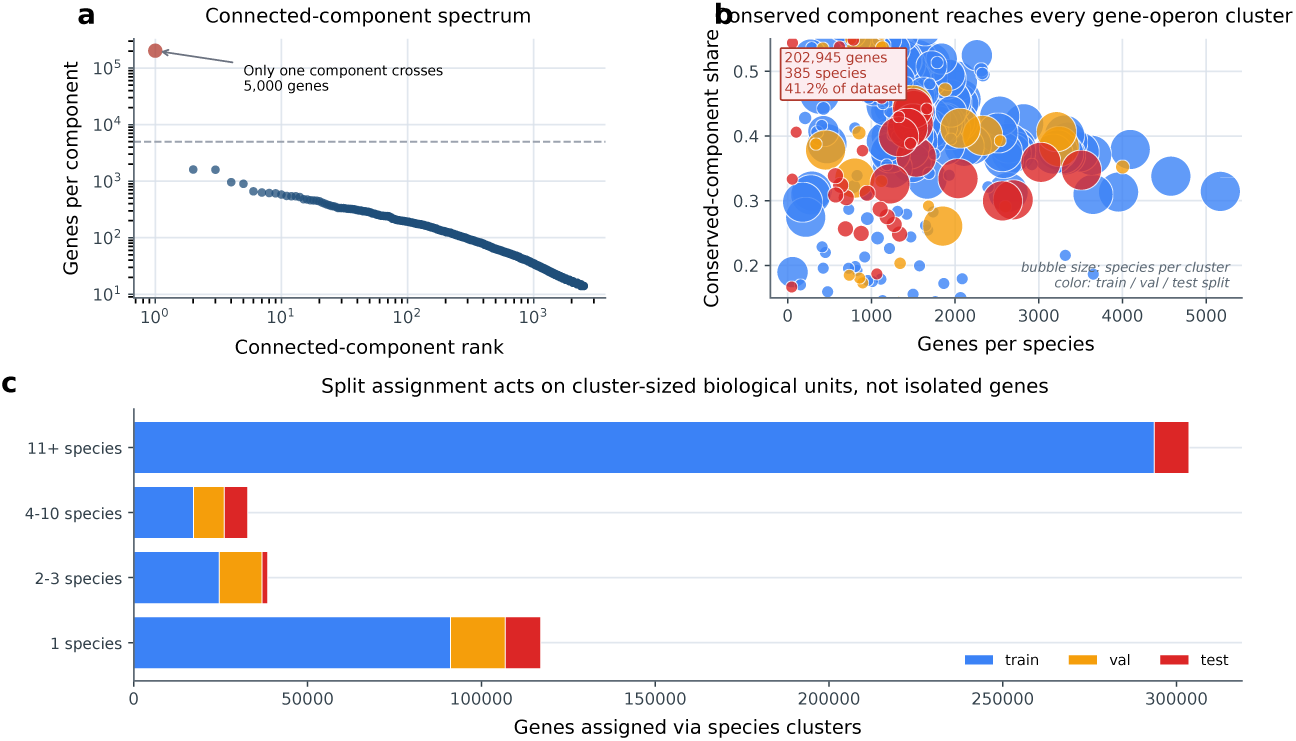
Leakage topology under the gene-operon split: conserved-component structure and cluster-level assignment. **a,** Connected-component spectrum over gene-operon clusters: one conserved component (202,945 genes, 41%) spans all 385 species; all others are *<*5,000 genes. **b,** Per-species conserved-component share (30–60%); the conserved component reaches every species (and therefore every gene-operon cluster that contains a conserved family), providing family-level shortcuts that inflate overall *ρ*. Bubble size encodes species-cluster size; color encodes train/val/test assignment. **c,** Gene-operon split assigns entire clusters (not genes) to train/val/test; the conserved component is distributed ∼80/10/10 across those splits.

**Fig. ED3.**
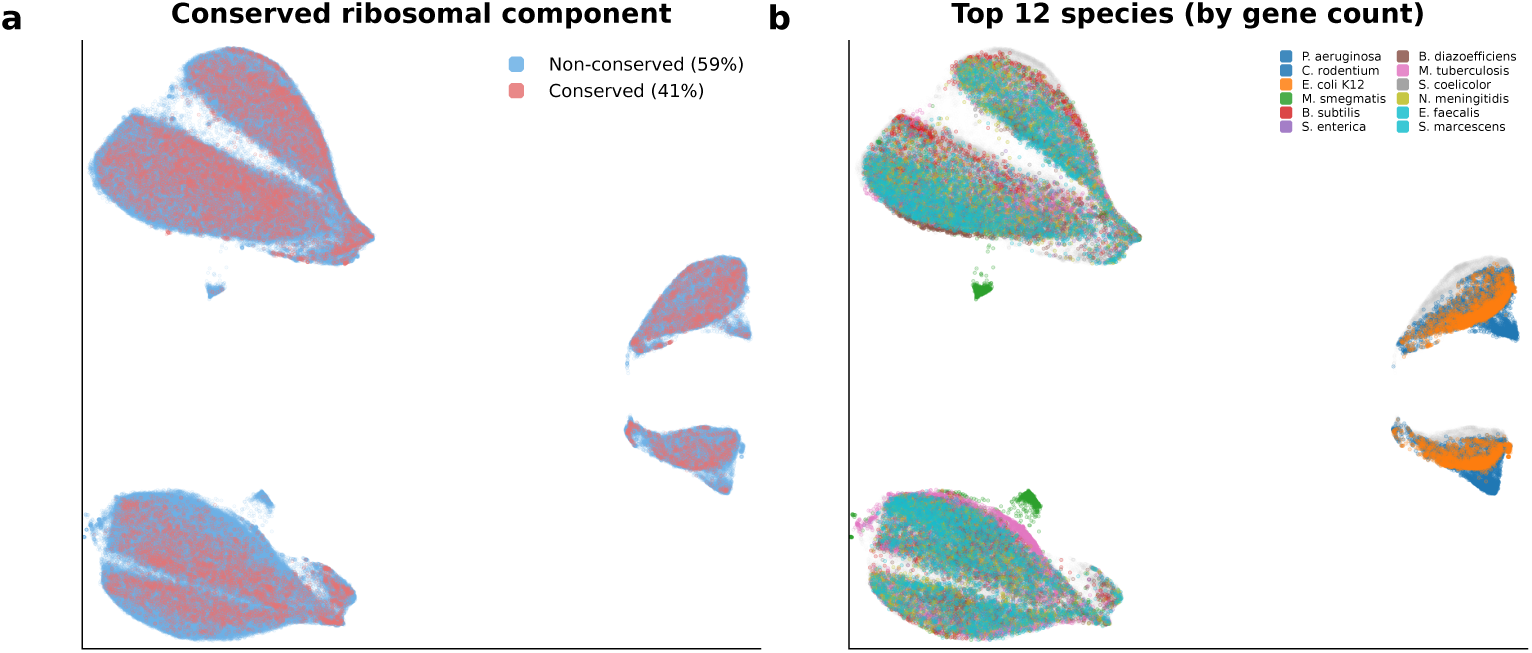
Anatomy of the gene expression manifold (XP5 fused representation, all 492,026 genes, UMAP). **a,** Conserved ribosomal component (red, 41%) occupies distinct spatial territory from non-conserved genes (blue, 59%). **b,** Top 12 species by gene count. Each species occupies a distinct spatial region; the expression-colored version of this manifold is in Fig. 2c.

## Supplementary Information

**Fig. S1.**
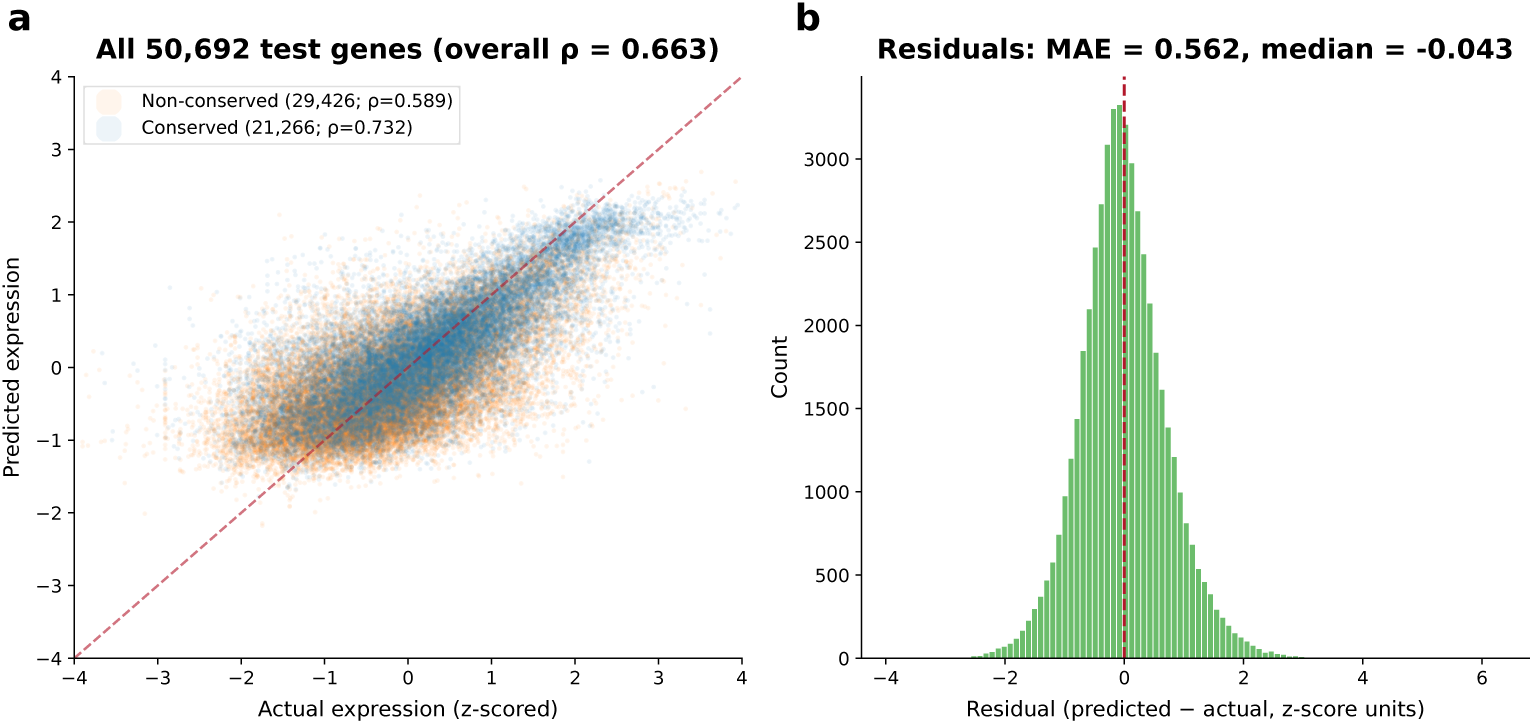
Prediction quality (XP5, gene-operon test set). **a,** Predicted versus actual expression for all 50,692 test genes. Conserved genes (blue; *ρ* = 0.732) and non-conserved genes (orange; *ρ*nc = 0.589) shown together; overall *ρ* = 0.663. **b,** Residual distribution (MAE in z-score units).

**Table S1.** Few-shot transfer to heterologous expression (Boël 2016, T7/BL21). Pretraining on native expression provides a warm-start advantage at *<*10% data but converges by 25%.

| Training fraction | Pretrained $\rho$ | Fresh ESM-C $\rho$ | $\Delta$ |
| --- | --- | --- | --- |
| 5% | $0.233 \pm 0.036$ | $0.196 \pm 0.057$ | +0.037 |
| 10% | $0.290 \pm 0.027$ | $0.261 \pm 0.021$ | +0.029 |
| 25% | $0.335 \pm 0.011$ | $0.328 \pm 0.012$ | +0.007 |
| 50% | $0.356 \pm 0.008$ | $0.359 \pm 0.007$ | -0.003 |
| 100% | $0.368 \pm 0.024$ | $0.362 \pm 0.038$ | +0.006 |

## Supplementary Note: Deployment ensemble breakdown

The Tier D_ensemble_ headline (*ρ*_nc_ = 0.606 ± 0.012) combines two independently motivated and standard ML ensembling forms: *retrain averaging* (different stochastic retrains of the same recipe) and *fusion-variant averaging* (different top-scoring fusion architectures with different modality compositions). Each is reported separately below and compared against the single-seed XP5 on the same five folds (rank-averaging throughout).

The two forms contribute nearly identical standalone gains (+0.012 and +0.013) and combine to +0.016, showing that they exploit partially orthogonal variance: retrain averaging smooths the stochastic-training component, while fusion-variant averaging mixes inherently different feature-use strategies. Neither form alone qualitatively changes the headline; the combined 5-member ensemble is reported in the main text only as a deployment convenience for practitioners who can afford 5× inference cost.

**Fig. S2.**
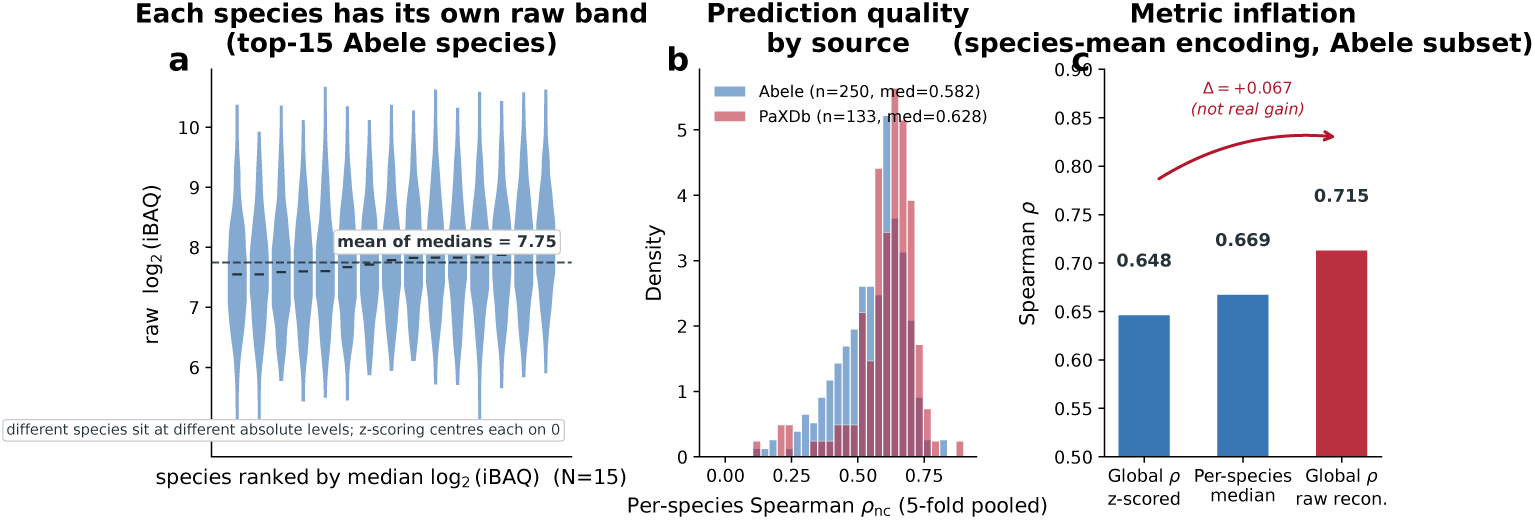
Why per-species z-scoring is necessary, and what it preserves. **a,** Top 15 Abele species by gene count, ranked by median raw log_2_(iBAQ). Each species occupies its own raw-abundance band (medians span ∼2 log_2_ units within this top-15 window, sitting inside the ∼5-orders-of-magnitude between-species variance over all 249 Abele species shown in Fig. 1f; within-species ranges 3–6 log_2_ units); per-species z-scoring centers each on 0 while leaving within-species rank correlations identical to those on the raw scale. **b,** Per-species *ρ*nc distribution by data source, computed by pooling each species’ predictions across the 5 gene-operon CV folds (one *ρ*nc per species; two small Abele species with *<*10 non-conserved test genes are excluded for *ρ*nc stability, *N* = 383 total). PaXDb (multi-study integrated, *n* = 133, median 0.628) outperforms Abele single-study iBAQ + small auxiliary depositions (*n* = 250, median 0.582); the ∼0.05 source gap is consistent with PaXDb’s cross-study noise averaging vs Abele’s single-study measurement noise. **c,** Apples-to-apples on the Abele test rows (the only subset with a defined raw log_2_ iBAQ scale): adding back per-species means inflates global *ρ* from 0.648 (z-scored) to 0.715 (raw-reconstructed; Δ*ρ* = +0.067). The species-mean component carrying this inflation is itself a measurement artifact (*ρ* = 0.23, *p* = 0.25 between PaXDb and Abele species means; *n* = 27 overlapping species). Z-scoring removes this confound; per-species medians (middle bar, 0.669) lie between the two globals.

All five ensemble members are seed-stable across independent retrains (two independent seeds, paired *t*-test on the same five folds): XP5 *p* = 0.34, Consensus 7-group *p* = 0.79, Central dogma *p* = 0.38 (Table S5).

**Fig. S3.**
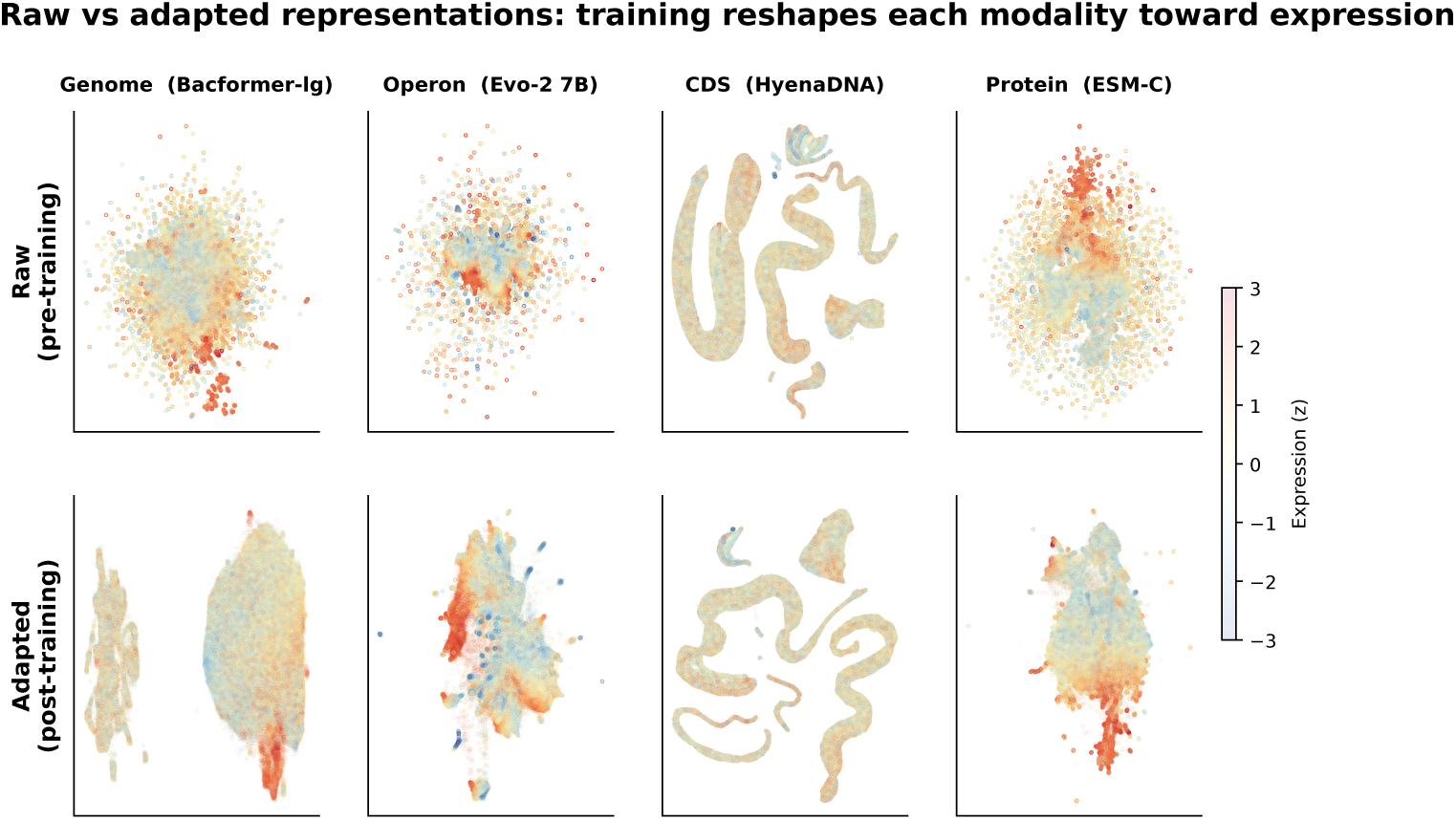
Training reshapes each modality toward expression (492,026 genes, colored by z-scored expression). **Top row:** Raw pre-trained embeddings. Each foundation model organizes genes by a different biological principle: chromosomal neighborhoods (Bacformer-large), operon-coherent clusters (Evo-2 7B), codon-composition gradients (HyenaDNA), protein-family clusters (ESM-C). No single raw embedding shows a clear expression gradient. **Bottom row:** Adapted embeddings (after per-modality pyramid MLP training). Each modality develops an expression-level gradient, though with different geometries. The fused representation (Fig. 2c; Extended Data Fig. ED3) integrates all four into a smooth manifold.

**Fig. S4.**
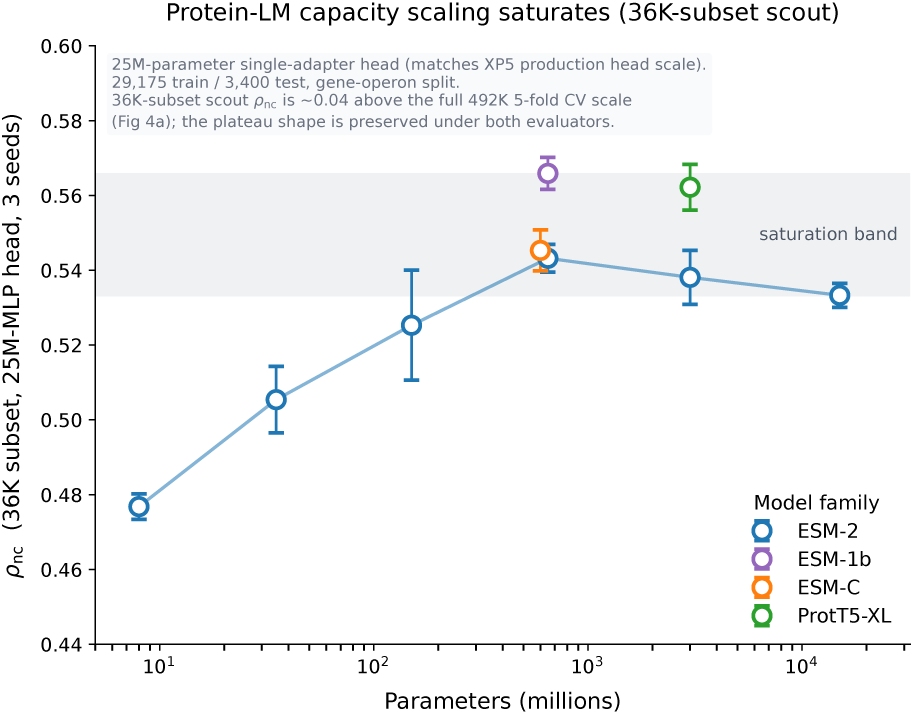
Protein-LM capacity scaling saturates across architectures (36K subset). Nine PLMs covering four families (ESM-2 from 8M to 15B, ESM-1b 650M, ESM-C 600M, and ProtT5-XL 3B), evaluated under the same single-adapter 25M-parameter MLP head used in the main XP5 pipeline. Fair-param head: hidden width binary-searched per model so every head has the same trainable budget; wider input embeddings do NOT get more head parameters, so ESM-2 15B (5120-d input) is not starved. 29,175 train / 3,670 val / 3,400 test (36,245-gene curated subset, gene-operon split; every point is an average over three independent random seeds, error bars = seed s.d.). ESM-2 rises sharply to 600M (*ρ*nc = 0.477 → 0.543 across 8M→650M) and then flattens or mildly declines: 3B gives 0.538, 15B gives 0.533. At matched head capacity ESM-2 15B does not exceed ESM-2 650M, so the plateau is not driven by an undersized head at this parameter count. ESM-C 600M (0.545), ESM-1b 650M (0.566), and ProtT5-XL 3B (0.562) cluster in the same plateau band despite different architectures and training objectives. *Test-set scale note.* This figure uses the curated 36K-subset 3,400-gene test (PaXDb-integrated multi-study labels). On the same 4 points with both evaluations, scout *ρ*nc on the 36K subset sits a systematic ∼0.04 above the full 492K 5-fold CV (main Fig. 4a). The plateau SHAPE is preserved under both evaluators; numbers on this SI figure are NOT directly comparable to XP5 *ρ*nc = 0.592 (Fig. 3), which is on the 492K scale. ESM-2 15B cannot be re-evaluated on the full 492K in the current manuscript cycle because its 5120-d embeddings are only extracted for the 36K curated subset.

**Fig. S5.**
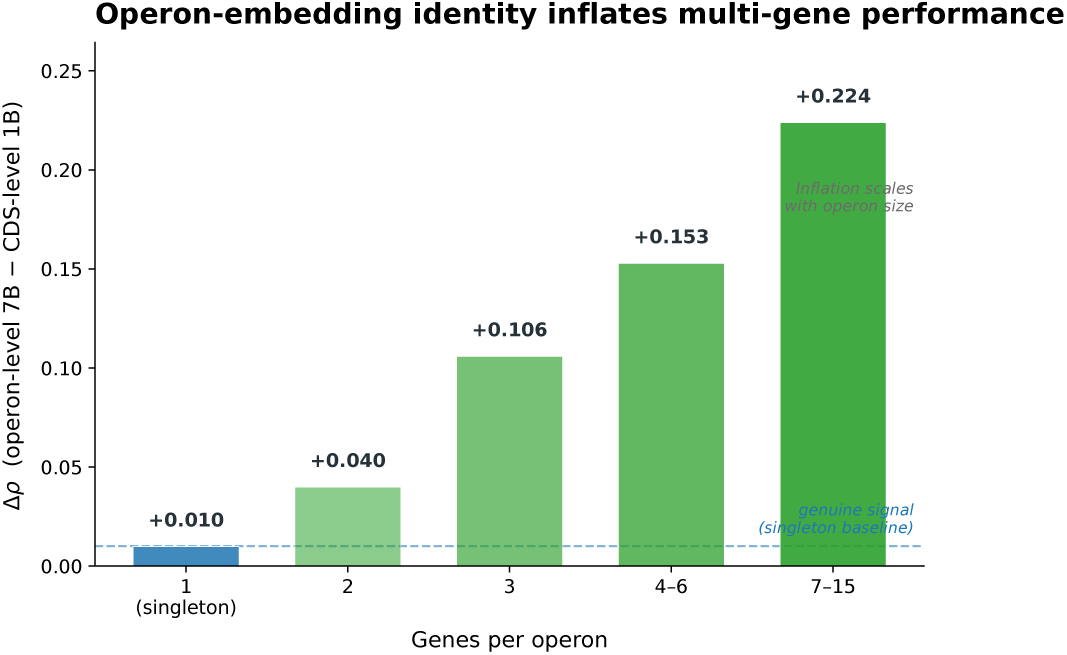
Operon-embedding identity inflates multi-gene performance. The operon-level encoder produces a single mean-pooled embedding for all genes in a polycistronic unit. When train and test genes share an operon, the model receives an identical input for both, an identity shortcut that inflates Δ*ρ*. Singleton genes (*N* = 35,644; leftmost bar) cannot leak and show the genuine DNA-scaling signal (Δ*ρ* = +0.010). Multi-gene operons show progressively larger inflation that scales with operon size, reaching Δ*ρ* = +0.224 for 7–15-gene operons. The headline Δ = +0.035 (all 50,692 test genes) is a mixture: +0.010 genuine (singletons) plus +0.101 leakage-inflated (multi-gene). Note: the 1B→7B comparison changes both model capacity and input window (60-nt init window for 1B vs full-operon mean-pool for 7B); the singleton Δ*ρ* = +0.010 reflects both factors.

**Fig. S6.**
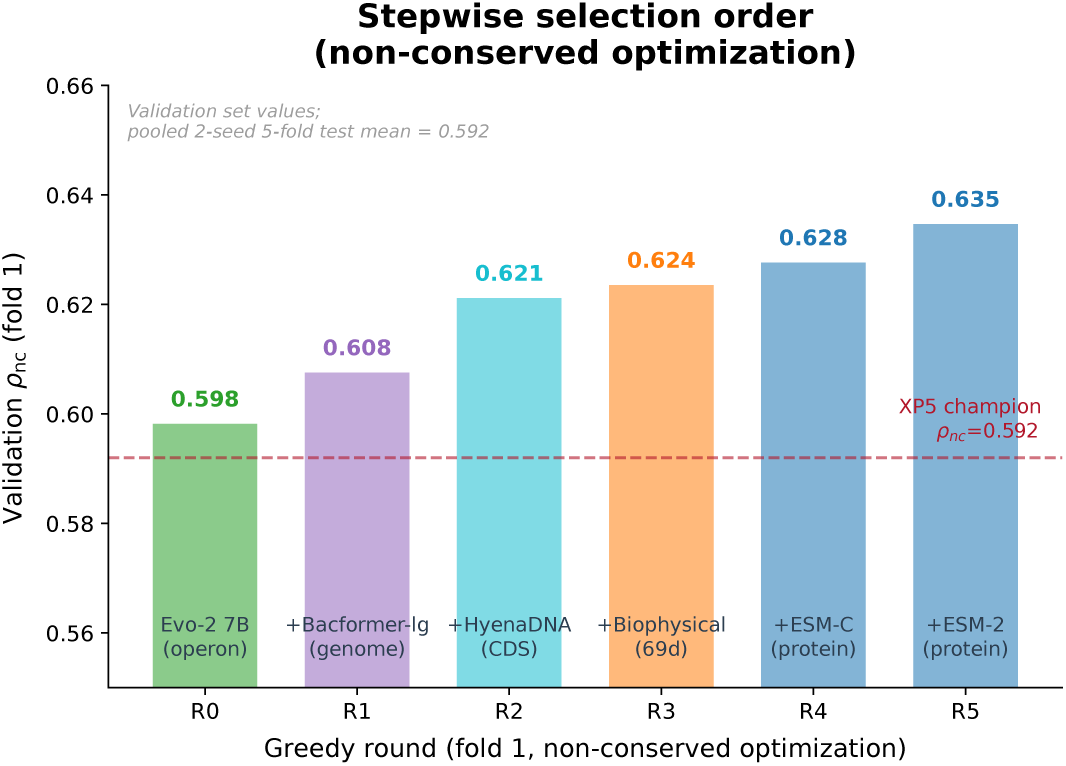
Greedy forward stepwise illustrates the selection order (fold 1, *ρ*nc optimization, validation set). Starting from Evo-2 7B (operon), each round adds the candidate that most improves validation *ρ*nc. DNA and genome-context modalities enter before protein: Bacformer-large (genome, R1), HyenaDNA (CDS, R2), biophysical features (R3), then ESM-C (protein, R4). Values shown are fold-1 validation *ρ*nc, which is higher than the pooled 2-seed five-fold test mean (0.592) due to fold-level variance. The selection order (not the absolute values) is the point: core modalities (operon + genome + CDS + biophysical) are selected across all folds; the protein model is the divergent 5th pick. Bar colors: green = operon, purple = genome, teal = CDS, orange = biophysical, blue = protein.

**Fig. S7.**
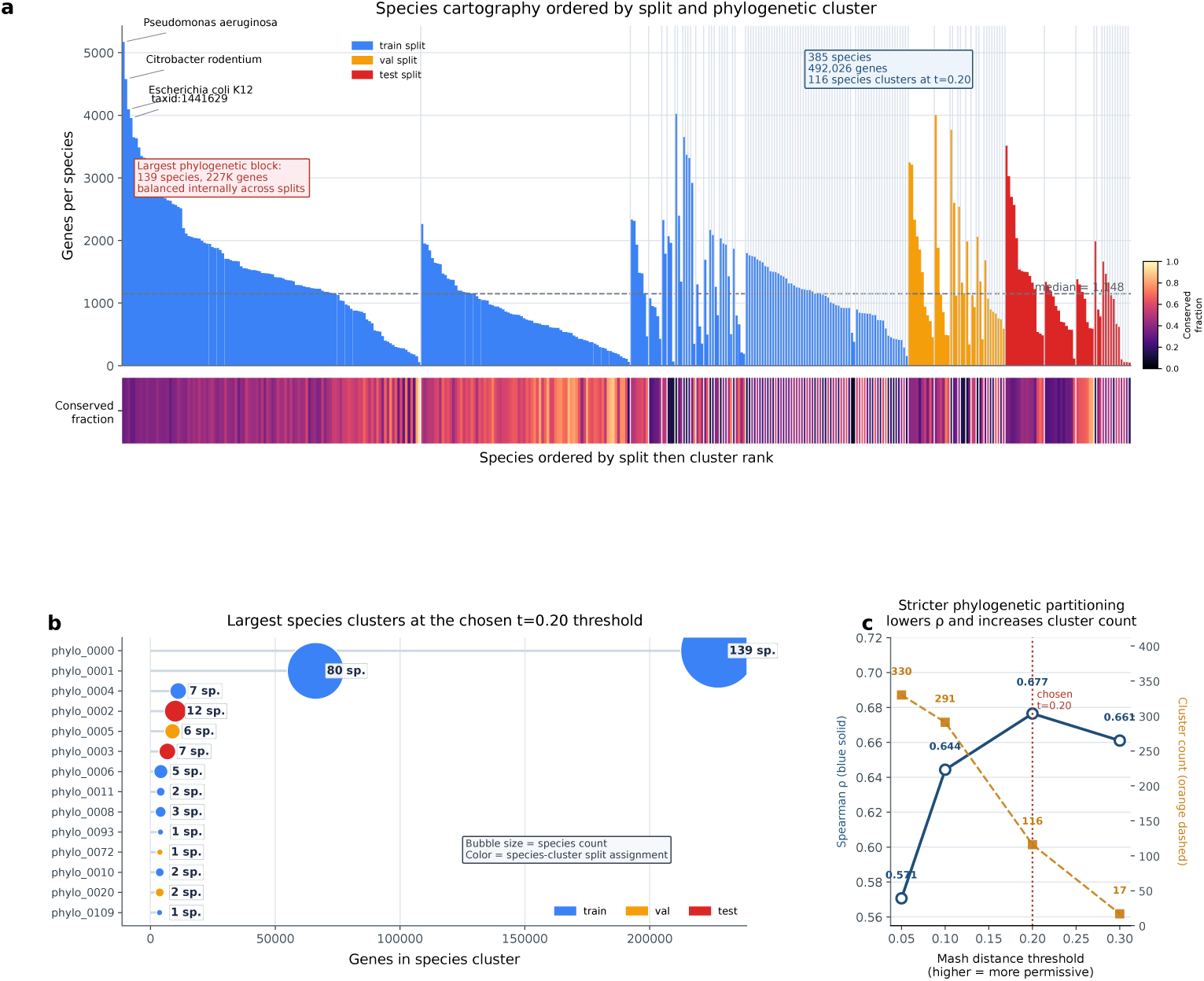
Species-cluster split structure and threshold dependence. **a,** Gene count per species, ordered by train/val/test assignment and then by species-cluster rank. The largest species cluster (139 species, 227K genes) is balanced internally across splits. Heatmap strip below shows the per-species conserved-ribosomal-component fraction. **b,** Largest species clusters at the chosen Mash *t* = 0.20 threshold. Bubble size encodes species count per cluster; bubble color encodes species-cluster split assignment. **c,** Stricter partitioning (lower Mash *t*) reduces overall *ρ* (blue, solid) and increases the number of phylogenetic clusters (orange, dashed) at the same 80/10/10 train/val/test proportions. Throughout the paper we use *species-cluster split* to refer to the single-linkage clustering at *t* = 0.20 that defines this partition, and *gene-operon split* to refer to the complementary protein-cluster split used as the primary evaluation regime.

**Fig. S8.**
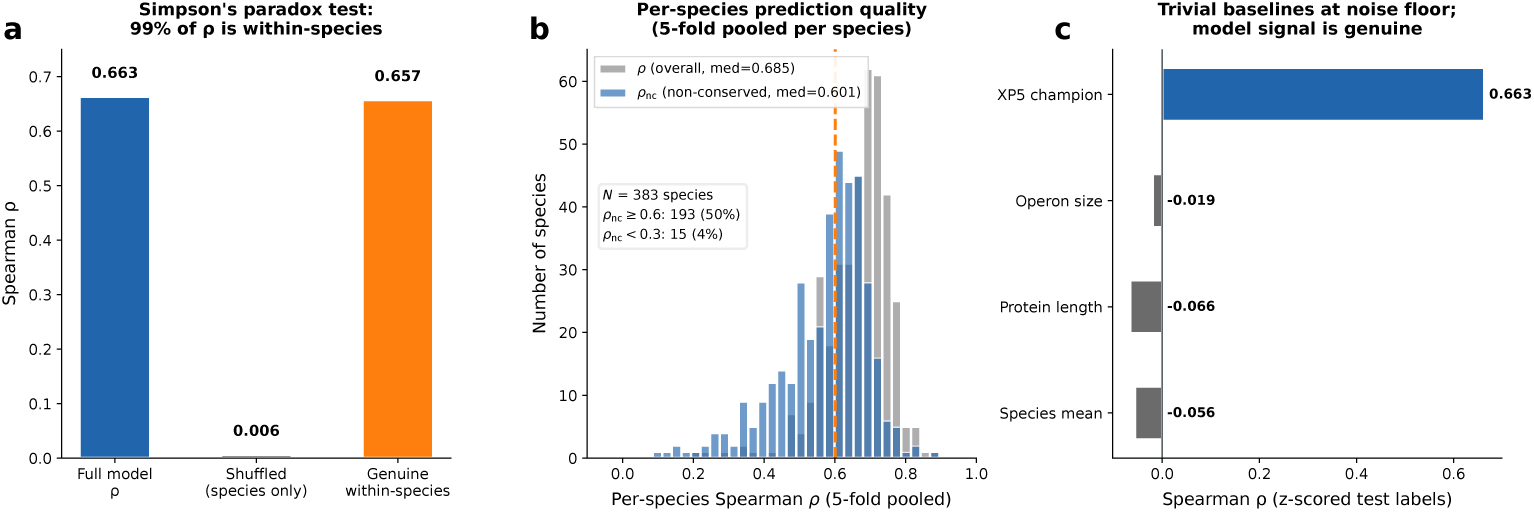
Expression confound analysis confirms genuine within-species prediction. **a,** Simpson’s paradox test: within-species label shuffling (destroys gene-level signal, preserves species means) reduces *ρ* from 0.663 to 0.006. XP5’s signal is 99% within-species. **b,** Per-species *ρ* distribution computed by pooling each species’ predictions across the 5 gene-operon CV folds (one *ρ* per species, *N* = 383 of 385 species; two small Abele species with *<*10 non-conserved test genes are excluded for *ρ*_nc_ stability). Median per-species *ρ*_overall_ = 0.685 vs pooled overall *ρ* = 0.663 (a +0.022 gap, the same family-recognition signal Simpson’s test removes); median per-species *ρ*nc = 0.601 vs pooled *ρ*nc = 0.589, a much smaller gap of +0.012, since the conserved-component contribution is excluded from *ρ*nc by construction. 50% of species reach *ρ*nc ≥ 0.6; 4% fall below 0.3 (the failure tail discussed in Methods). **c,** Trivial baselines (species mean, protein length, operon size) are all at noise floor (|*ρ*| *<* 0.07), confirming XP5’s *ρ* = 0.663 is genuine learned signal.

**Table S2.** Recipe comparison (five-fold CV, gene-operon splits, representative subset). Top rows: fusion variants around XP5. Middle: representative R1 pairs showing partner interchangeability. Bottom: scale-ablation baselines and deployment-tier recipes. All recipes: same five Union-Find zero-leakage folds with non-conserved checkpoint selection; single random seed unless noted; *ρ*_nc_ = non-conserved Spearman *ρ*; mod groups in brackets: G=genome, O=operon, C=CDS, P=protein, B=biophysical.

| Recipe | Mods | Scales | $\rho_{ov}$ | $\rho_{nc}$ |
| --- | --- | --- | --- | --- |
| *XP5 and near-XP5 variants* |  |  |  |  |
| Consensus (7-group union) | 7 | GOCPB | $0.670 \pm 0.013$ | **$0.594 \pm 0.011$** |
| **XP5** (pooled 2-seed) | 5 | GOCPB | $0.667 \pm 0.014$ | $0.592 \pm 0.011$ |
| XP5 + dropout ($p=0.2$) | 5 | GOCPB | $0.665 \pm 0.009$ | $0.590 \pm 0.014$ |
| Central dogma (BioLangFusion analogue) | 4 | OCP | $0.662 \pm 0.011$ | $0.586 \pm 0.012$ |
| All 17 modalities | 17 | GOCPB | $0.659 \pm 0.009$ | $0.583 \pm 0.009$ |
| *Evo-2 7B + one partner (all within $\Delta\rho_{nc} = 0.006$)* |  |  |  |  |
| + ProtT5-XL | 2 | OP | $0.648 \pm 0.012$ | $0.575 \pm 0.011$ |
| + ESM-C | 2 | OP | $0.651 \pm 0.012$ | $0.574 \pm 0.009$ |
| + HyenaDNA | 2 | OC | $0.648 \pm 0.011$ | $0.573 \pm 0.010$ |
| + CodonFM | 2 | OC | $0.647 \pm 0.011$ | $0.569 \pm 0.010$ |
| *Scale ablations and single-modality baselines* |  |  |  |  |
| Evo-2 7B solo | 1 | O | $0.644 \pm 0.015$ | $0.565 \pm 0.011$ |
| All DNA (6 models) | 6 | GOCB | $0.652 \pm 0.016$ | $0.573 \pm 0.013$ |
| XP5 – Evo-2 7B | 4 | GCPB | $0.628 \pm 0.011$ | $0.563 \pm 0.015$ |
| ESM-C + ProtT5-XL (Tier A) | 2 | P | $0.579 \pm 0.011$ | $0.518 \pm 0.011$ |
| ESM-C solo | 1 | P | $0.575 \pm 0.013$ | $0.509 \pm 0.013$ |
| BF-large solo | 1 | G | $0.545 \pm 0.012$ | $0.475 \pm 0.010$ |
| All RNA (2 models) | 2 | — | $0.199 \pm 0.016$ | $0.153 \pm 0.008$ |

Biophysical = 1 modality group (5 feature blocks). Full 32-recipe table available in the repository; the entries shown here cover every scale ablation and every locked deployment tier discussed in Table ED4.

**Table S3.** Effect of removing Aiki-XP training-set near-neighbors from external benchmarks. For each benchmark, proteins sharing ≥30% sequence identity and ≥80% query coverage with the Aiki-XP 492K training set were removed (MMseqs2 easy-search). This controls for homology leakage in *our* evaluation; it does not imply any train/test overlap issue in NetSolP or MPEPE themselves, since neither was trained on our data. Δ*ρ* = *ρ*_clean_ − *ρ*_full_; significance (∗∗) marks rows whose 95% bootstrap CI excludes zero (*n* = 1,000 resamples). The key finding for Aiki-XP is that protein-only tiers (Tier A, ESM-C solo) are robust to near-neighbor removal (|Δ*ρ*| *<* 0.01), while the legacy F10 DNA-based multimodal recipe collapses (Δ*ρ* = −0.073 on Boël), indicating that DNA-modality features can inflate performance via homolog recognition on heterologous benchmarks. Tool citations: NetSolP[21], MPEPE[20], CaLM[32]; XP5 rows are this work.

| Tool | Benchmark | $N_{\text{full}}$ | $N_{\text{clean}}$ | $\rho_{\text{full}}$ | $\rho_{\text{clean}}$ | $\Delta\rho$ | Sig. |
| --- | --- | --- | --- | --- | --- | --- | --- |
| *Boël 2016 (T7, E. coli; 61% of proteins share $\geq 30\%$ identity with Aiki-XP training set)* |  |  |  |  |  |  |  |
| NetSolP usability | Boël | 6,348 | 2,481 | 0.342 | 0.303 | **-0.038** | ** |
| NetSolP solubility | Boël | 6,348 | 2,481 | 0.135 | 0.153 | +0.018 |  |
| MPEPE | Boël | 6,348 | 2,481 | 0.565 | 0.567 | +0.001 |  |
| ESM-C solo | Boël | 6,348 | 2,481 | 0.192 | 0.190 | -0.002 |  |
| Tier A | Boël | 6,348 | 2,481 | 0.165 | 0.172 | +0.008 |  |
| XP5 classical (69d) | Boël | 6,348 | 2,481 | 0.159 | 0.148 | -0.011 |  |
| CaLM (Ridge 20K) | Boël | 6,348 | 2,481 | 0.146 | 0.135 | -0.011 |  |
| F10 (legacy multimodal) | Boël | 6,348 | 2,481 | 0.078 | 0.004 | **-0.073** | ** |
| *Price/NESG 2011 (cell-free; 66% overlap)* |  |  |  |  |  |  |  |
| NetSolP usability | Price/NESG | 9,703 | 3,299 | 0.329 | 0.212 | **-0.117** | ** |
| NetSolP solubility | Price/NESG | 9,703 | 3,299 | 0.040 | -0.038 | **-0.078** | ** |
| MPEPE | Price/NESG | 9,703 | 3,299 | 0.312 | 0.222 | **-0.090** | ** |
| ESM-C solo | Price/NESG | 9,703 | 3,299 | 0.137 | 0.114 | -0.023 |  |
| Tier A | Price/NESG | 9,703 | 3,299 | 0.113 | 0.103 | -0.010 |  |
| XP5 classical (69d) | Price/NESG | 9,703 | 3,299 | 0.117 | 0.127 | +0.011 |  |
| CaLM (Ridge 20K) | Price/NESG | 9,703 | 3,299 | 0.101 | 0.080 | -0.021 |  |
| F10 (legacy multimodal) | Price/NESG | 9,703 | 3,299 | 0.049 | 0.070 | +0.021 |  |
| *SSGCID (heterologous structural bio.; 70% overlap)* |  |  |  |  |  |  |  |
| MPEPE | SSGCID | 14,507 | 4,364 | 0.095 | 0.171 | **+0.077** | ** |
| Tier A | SSGCID | 14,507 | 4,364 | 0.061 | 0.073 | +0.013 |  |
| *Synechococcus elongatus (native DIA-MS; 68% overlap)* |  |  |  |  |  |  |  |
| MPEPE | Synechococcus | 2,018 | 642 | -0.031 | -0.000 | +0.031 |  |

**Table S4.** Decomposition of the Tier D_ensemble_ advantage into retrain-and fusion-variant averaging. Each group is a rank-average ensemble evaluated on the same five gene-operon folds; Δ and *p* are paired per-fold against the single-seed XP5 baseline. Retrain members: XP5 at two independent seeds + an XP5 variant trained with modality dropout. Fusion-variant members: XP5 + Consensus 7-group + Central dogma (BioLangFusion analogue). Combined: union of both groups (5 members total).

| Group | $\rho_{\text{nc}}$ | $\Delta$ | $p$ | Wins |
| --- | --- | --- | --- | --- |
| XP5 (single-seed baseline) | $0.590 \pm 0.014$ | — | — | — |
| Retrain-avg (3 members) | $0.602 \pm 0.012$ | +0.012 | 0.0003 | 5/5 |
| Fusion-variant avg (3 members) | $0.603 \pm 0.013$ | +0.013 | 0.0006 | 5/5 |
| **5-member combined** | **$0.606 \pm 0.012$** | **+0.016** | **0.0002** | **5/5** |

**Table S5.** Seed stability of all ensemble members. Each recipe trained at two independent seeds on the same five gene-operon folds; paired *t*-test on per-fold *ρ*nc.

| Recipe | Seed 42 $\rho_{\text{nc}}$ | Seed 0 $\rho_{\text{nc}}$ | $\Delta$ | $p$ |
| --- | --- | --- | --- | --- |
| XP5 | $0.590 \pm 0.014$ | $0.593 \pm 0.010$ | +0.002 | 0.34 |
| XP5 + mod-dropout | single-seed (retrain variant; stability inherited from XP5) |  |  |  |
| Consensus 7-group | $0.594 \pm 0.012$ | $0.594 \pm 0.016$ | −0.001 | 0.79 |
| Central dogma | $0.586 \pm 0.014$ | $0.588 \pm 0.010$ | +0.003 | 0.38 |

## References

[1] Szymborski, J. & Emad, A. A flaw in using pretrained protein language models in protein–protein interaction inference models. Nat. Mach. Intell. 8, 197–208 (2026).

[2] Rafi, A.M., Kiyota, B., Yachie, N. & de Boer, C.G. Detecting and avoiding homology-based data leakage in genome-trained sequence models. bioRxiv 2025.01.22.634321 (2025).

[3] Hallee, L., Peleg, T., Rafailidis, N. & Gleghorn, J.P. Protein Language Models are Accidental Taxonomists. bioRxiv 2025.10.07.681002 (2025).

[4] Huang, Q., Szklarczyk, D., Oehninger, J. & von Mering, C. PaxDb v6.0: reprocessed, LLM-selected, curated protein abundance data across organisms. Nucleic Acids Res. 54, D427–D439 (2026).

[5] Abele, M. et al. Proteomic diversity in bacteria: insights and implications for bacterial identification. Mol. Cell. Proteomics 24, 100917 (2025).

[6] Lathe, W.C., Snel, B. & Bork, P. Gene context conservation of a higher order than operons. Trends Biochem. Sci. 25, 474–479 (2000).

[7] Lin, Z. et al. Evolutionary-scale prediction of atomic-level protein structure with a language model. Science 379, 1123–1130 (2023).

[8] Nguyen, E. et al. HyenaDNA: long-range genomic sequence modeling at single nucleotide resolution. Adv. Neural Inf. Process. Syst. 36 (2023).

[9] Zhou, Z. et al. DNABERT-2: efficient foundation model and benchmark for multi-species genome. ICLR (2024).

[10] Dalla-Torre, H. et al. The Nucleotide Transformer: building and evaluating robust foundation models for human genomics. Nat. Methods 22, 287–297 (2025).

[11] Penić, R.J., Vlašić, T., Huber, R.G., Wan, Y. & Šikić, M. RiNALMo: general-purpose RNA language models can generalize well on structure prediction tasks. Nat. Commun. 16, 5671 (2025).

[12] Brixi, G. et al. Genome modeling and design across all domains of life with Evo 2. bioRxiv (2025).

[13] Wiatrak, M. et al. A contextualised protein language model reveals the functional syntax of bacterial evolution. bioRxiv 2025.07.20.665723 (2025).

[14] Sharp, P.M. & Li, W.H. The codon adaptation index, a measure of directional synonymous codon usage bias. Nucleic Acids Res. 15, 1281–1295 (1987).

[15] Lorenz, R. et al. ViennaRNA Package 2.0. Algorithms Mol. Biol. 6, 26 (2011).

[16] Salis, H.M., Mirsky, E.A. & Voigt, C.A. Automated design of synthetic ribosome binding sites to control protein expression. Nat. Biotechnol. 27, 946–950 (2009).

[17] Kudla, G., Murray, A.W., Tollervey, D. & Plotkin, J.B. Coding-sequence determinants of gene expression in *Escherichia coli*. Science 324, 255–258 (2009).

[18] Steinegger, M. & Söding, J. MMseqs2 enables sensitive protein sequence searching for the analysis of massive data sets. Nat. Biotechnol. 35, 1026–1028 (2017).

[19] Fu, G., Yan, Y., Chen, Y. & Shao, B. Predicting microbial transcriptome using genome sequence. bioRxiv 2024.12.30.630741 (2024).

[20] Ding, Z. et al. MPEPE, a predictive approach to improve protein expression in *E. coli* based on deep learning. Comput. Struct. Biotechnol. J. 20, 1142–1153 (2022).

[21] Thumuluri, V., Martiny, H.-M., Almagro Armenteros, J.J., Salomon, J., Nielsen, H. & Johansen, A.R. NetSolP: predicting protein solubility in *Escherichia coli* using language models. Bioinformatics 38, 941–946 (2022).

[22] Buric, F. et al. Amino acid sequence encodes protein abundance shaped by protein stability at reduced synthesis cost. Protein Sci. 34, e5239 (2025).

[23] Russo, D.A., Schneidmadel, F.R. & Zedler, J.A.Z. Library-free data-independent acquisition mass spectrometry enables comprehensive coverage of the cyanobac-terial proteome. Plant Physiol. 199, kiaf334 (2025).

[24] Li, G.-W., Burkhardt, D., Gross, C. & Weissman, J.S. Quantifying absolute protein synthesis rates reveals principles underlying allocation of cellular resources. Cell 157, 624–635 (2014).

[25] Mori, M. et al. From coarse to fine: the absolute *Escherichia coli* proteome under diverse growth conditions. Mol. Syst. Biol. 17, e9536 (2021).

[26] Taniguchi, Y. et al. Quantifying *E. coli* proteome and transcriptome with single-molecule sensitivity in single cells. Science 329, 533–538 (2010).

[27] Cambray, G., Guimaraes, J.C. & Arkin, A.P. Evaluation of 244,000 synthetic sequences reveals design principles to optimize translation in *Escherichia coli*. Nat. Biotechnol. 36, 1005–1015 (2018).

[28] Klumpp, S., Scott, M., Pedersen, S. & Hwa, T. Molecular crowding limits translation and cell growth. Proc. Natl Acad. Sci. USA 110, 16754–16759 (2013).

[29] Mollaysa, A. et al. BioLangFusion: multimodal fusion of DNA, mRNA, and protein language models. ICML 2025 Workshop on Multi-modal Foundation Models and Large Language Models for Life Sciences (2025).

[30] Zhang, Y. et al. Predicting functions of uncharacterized gene products from microbial communities. Nat. Biotechnol. (2025). 10.1038/s41587-025-02813-7.

[31] Ellington, C.N. et al. Rapid and reproducible multimodal biological foundation model development with AIDO.ModelGenerator. ICML 2025 Workshop on Generative AI and Biology (2025).

[32] Outeiral, C. & Deane, C.M. Codon language embeddings provide strong signals for use in protein engineering. Nat. Mach. Intell. 6, 170–179 (2024).

[33] Tankhilevich, E. et al. RP3Net: a deep learning model for predicting recombinant protein production in *Escherichia coli*. Bioinformatics 42, btag003 (2026).

[34] Shen, Y., Kudla, G. & Oyarzun, D.A. Improving the generalization of protein expression models with mechanistic sequence information. Nucleic Acids Res. 53, gkaf020 (2025).

[35] Elnaggar, A. et al. ProtTrans: toward understanding the language of life through self-supervised learning. IEEE Trans. Pattern Anal. Mach. Intell. 44, 7112–7127 (2022).

[36] Heuschkel, J., Kingsley, L.J., Pefaur, N., Nixon, A. & Cramer, S. Advancing codon language modeling with synonymous codon constrained masking. Nucleic Acids Res. 54, gkag166 (2026).

[37] Nikolados, E.-M., Wongprommoon, A., Mac Aodha, O., Cambray, G. & Oyarzun, D.A. Accuracy and data efficiency in deep learning models of protein expression. Nat. Commun. 13, 7755 (2022).

[38] James, T., Williamson, B., Tino, P. & Wheeler, N. Whole-genome phenotype prediction with machine learning: open problems in bacterial genomics. Bioinformatics 41, btaf206 (2025).

[39] Rao, V.M. et al. Generalist biological artificial intelligence in modeling the language of life. Nat. Biotechnol. (2026).

[40] Nieuwkoop, T., Finger-Bou, M., van der Oost, J. & Claassens, N.J. The ongoing quest to crack the genetic code for protein production. Mol. Cell 80, 193–209 (2020).

[41] Calero, P. & Nikel, P.I. Chasing bacterial chassis for metabolic engineering: a perspective review from classical to non-traditional microorganisms. Microb. Biotechnol. 12, 98–124 (2019).

[42] Morris, R., Black, K.A. & Stollar, E.J. Uncovering protein function: from classification to complexes. Essays Biochem. 66, 255–285 (2022).

[43] Gonlepa, M.K., Osotuyi, T.B., Ofuonye, C.G. & Durojaye, O.A. Protein engineering as a driver of innovation in therapeutics biotechnology and the global bioeconomy. Discov. Chem. 2, 226 (2025).

[44] Jacob, F. & Monod, J. Genetic regulatory mechanisms in the synthesis of proteins. J. Mol. Biol. 3, 318–356 (1961).

